# Causal inference for multiple risk factors and diseases from genomics data

**DOI:** 10.1101/2023.12.06.570392

**Authors:** Nick Machnik, Mahdi Mahmoudi, Malgorzata Borczyk, Ilse Krätschmer, Markus J. Bauer, Matthew R. Robinson

## Abstract

Statistical causal learning in genome-wide association studies (GWAS) relies on the instrumental variable method of Mendelian Randomization (MR). Currently, an overwhelming number of MR studies purport to show causal relationships among a wide range of risk factors and outcomes. Here, we find that naive application of many recently proposed MR approaches results in numerous null relationships being discovered as highly significant. We show that a well-controlled error rate can be achieved through a graphical inference approach which: (i) selects a set of genetic instrumental variables (IVs) from GWAS summary static controlling for LD, linkage and pleiotropy; (ii) accommodates rare variants and binary outcomes in a principled way; (iii) distinguishes direct from indirect risk factors in very high-dimensional data; and (iv) identifies potential unobserved latent confounding. Only 20 minutes of wall-clock compute time is required for our Causal Inference GWAS (CI-GWAS) approach to jointly analyze a set of 9 common health risk factors, four common complex metabolic disease outcomes and 8.4M genetic variants recorded for 458,747 individuals in the UK Biobank. Genome-wide, we find that very few genetic variants are suitable MR IVs, with only 696 variants remaining when analyzing all traits jointly. While we replicate almost all paths previously found by CAUSE between risk factors and outcomes, we show that only few of these reflect direct adjacencies and that many cannot be distinguished from unmeasured confounding within the UK Biobank data. Our results suggest that well-curated longitudinal records and family data are likely needed to overcome the mixtures of temporal precedence and reverse-causality in biobank data. Our approach provides a first-step toward robust principled screening for potential causal links to understand the underlying nature of phenotypic correlations in biobank data.

## Introduction

Distinguishing causation from correlation is fundamentally important for understanding the etiology of common complex disease. Randomized controlled trials (RCTs) are the gold standard for causal inference, but their difficulty, expense, and often also ethical considerations, prohibit testing for potential disease-associated risk factors among the vast array of human complex traits. As a result, the most commonly used methods for statistical causal inference in a genetic or biomedical setting center around the instrument variable (IV) framework of Mendelian randomization (MR) [1, 2]. A wide variety of MR approaches have been developed to test whether there exists a potential causal effect, direct or indirect, of one or more mediator variable(s) onto one or more outcome variable(s) [3, 4, 5, 6, 7, 8, 9, 10, 11, 12, 13, 14, 15]. These have been applied in an overwhelming number of recent MR studies that all purport to show causal relationships among a wide range of risk factors and outcomes. However, while easy to apply in practice, there are major limitations to MR that may make inference unreliable: *(i)* testing is conducted using a single form of fixed graphical pattern; *(ii)* SNPs with strongest genotype-phenotype association test statistics are used as IVs, which relies on a number of key assumptions that are difficult to explicitly test; and *(iii)* the type-I error of the significance testing may be very high [16].

Theoretical developments in statistical causal inference show that graphical modeling can be used to learn the partial correlations among hundreds of variables, whilst controlling for potential unobserved confounding within a single model [17, 18, 19]. However, there are currently no algorithms capable of conducting this form of causal discovery at scale. Developing these approaches to accommodate highly correlated variables of various distributions and millions of highly correlated genetic markers, whilst allowing data to be combined across biobanks, would facilitate accurate high-throughput screening of potential causal links among modifiable risk factors and disease outcomes.

Here, we propose an approach for conducting Causal Inference in Genome-Wide Association Studies (CI-GWAS), that is capable of inferring a single graph at scale describing the relationships among millions of genetic markers, tens of risk factors and multiple disease outcomes. We utilise the inferred graph for more principled detection of potential IVs, controlling for linkage disequilibrium genome-wide and fulfilling the core MR assumptions, in particular the absence of horizontal pleiotropy and potential latent confounding of the marker outcome relationship. In simulations, we show that CI-GWAS selects appropriate IVs and that this improves upon the high false discovery rates of current MR approaches. We show that this holds even when using rare variants and rare disease outcomes when placing relationships on the liability-scale to correct for differences in outcome prevalence. Applying CI-GWAS to summary statistic data of 8.4M SNPs and 15 phenotypes requires only 20 minutes of wall-clock compute time on a standard GPU server. We find that few GWAS variants are suitable MR IVs, but that almost all previously reported paths among risk factors and outcomes [11] can be recovered. With further investigation, we find that almost all these risk factor-outcome paths reflect either an indirect path through another trait or unmeasured confounding. Our approach can be used on individual-level or summary statistic data and provides a solution to facilitate robust one- or two-sample MR inference on large-scale genotype-phenotype data.

## Results

### The structure of CI-GWAS

CI-GWAS is a method that aims to learn a graphical model that describes the causal relationships among genetic and phenotypic variables. It consists of two parts: Part 1 is the construction of a skeleton, an undirected graph that represents the possible direct causal relationships between all observed variables in the model; Part 2 then directs the edges in the skeleton to obtain a partial ancestral graph (PAG), while taking into account potential confounding through latent variables. The graph learned by CI-GWAS can then be used in conjunction with an MR method to infer a set of potential causal relationships and their effects.

As we describe in the Methods and show in simulations below, part 1 identifies risk factors that are most likely to have direct effects on (a partial correlation with) outcomes and yields valid genetic IVs for multivariate MR. Our algorithm, based on a GPU-accelerated version of the PC-stable algorithm [20, 21], is additionally accelerated by iterative variable selection and by exploiting the LD-block structure of the genome. This enables us to construct a complete marker-trait graph, for millions of markers and tens of traits, despite the exponential run-time scaling in the number of variables. This extends previously proposed graphical methods that were limited to few hundreds [22] or thousands [23] of genetic variables or relied on the construction of separate graphs for trait-trait and marker-trait relationships with pre-selected genetic variants by GWAS [16].

Part 2 is based on the RFCI algorithm [19]. We introduce a novel way to obtain order-independent separation sets that offer a conservative way of orienting trait relationships, that is, to learn which phenotypes have evidence for a direct relationship with each outcome. We also incorporate prior knowledge that markers precede traits temporally, which dramatically reduces computation time. We denote our version of RFCI as sRFCI.

Combining graphical inference with standard MR methods takes advantage of the principled instrumental selection in part 1. This also advances previous work on graphical inference [16] because now marker selection is made through joint estimation, where many variables can be conditioned on at once, and pleiotropic markers can be used for inference, promising more accurate IV selection and improved inference in MR as shown below.

CI-GWAS takes as input (i) the matrix of marker-maker correlations (LD matrix); (ii) a matrix of marker-trait correlations; (iii) a matrix of trait-trait correlations. This makes CI-GWAS applicable to summary statistic data, as LD-matrices, effect sizes of marker-trait associations, minor-allele frequencies, trait variances and trait-trait correlations can be shared whilst preserving privacy.

Part 1 of our algorithm relies on partial correlations vanishing for (conditionally) independent variables, which generally holds for Gaussian variables [24] but not necessarily for binary or ordinal variables. We find that treating binary case-control outcomes like Gaussian variables can lead to largely elevated numbers of false positive findings, especially for independence tests between markers with very low minor-allele-frequencies and binary traits with very unbalanced numbers of cases and controls (Figure S1). As has been shown in previous work [25], a heterologous correlation matrix with Pearson correlations between Gaussian, polyserial between Gaussian and ordinal and polychoric correlations between ordinal variables can be used to prevent this issue. This effectively places binary variables on the liability scale and has the additional advantage of allowing comparisons among traits at the same scale accounting for differences in outcome prevalence. At the sample size lowest minor-allele-frequency and lowest trait prevalence in the UK Biobank data used in this study, using polyserial correlations between markers and case-control traits is effective in controlling the numbers of false positives (Figure S1).

### Reducing false positives in Mendelian Randomization

We first demonstrate that the selection of appropriate IVs can reduce the number of false positive discoveries in MR. We conducted a simple simulation study based on a structural equation model consisting of two continuous traits *A* and *B* each with a set of 200 causal markers *X_A_* and *X_B_*chosen at random from 654,172 imputed SNP markers on chromosome 1 recorded for 458,746 individuals in the UKB. We allow for some shared fraction between *X_A_*and *X_B_* of size *γ*|*X_A_*|. We simulate data for *A* and *B* using causal effect sizes *β* ∈ {0, 0.25} for *A* → *B*, with marker effects chosen such that the heritability of each trait is fixed at 0.2. We then ran inverse variance weighting MR (IVW) with three different sets of IVs: (i) the parent markers selected with CI-GWAS; (ii) the true causal markers selected to simulate the traits (oracle); and (iii) markers with marginally significant effects at the genome-wide level (*α <* 10^−8^), identified in a GWAS (glm + clump, i.e. just selecting the top associated variants). We find that in the absence of a true causal effect between *A* and *B*, the approach with GWAS selected IVs tends to find spurious associations at a frequency that increases with the p-value threshold, while IVW run with oracle or CI-GWAS selected markers returns almost no false positive associations irrespective of the cutoff (Figure 1a). Only at a large cutoff point of *α <* 10^−1^, does the false positive rate increase for both the oracle and CI-GWAS IV selection. In the presence of a true causal effect between *A* and *B*, both the markers identified by CI-GWAS, as well as the oracle markers, result in causal link FPRs below or at 0.05 up to a p-value threshold of *α <* 10^−2^ (Figure 1b). In contrast, GWAS-selected IVs give a much higher FPR at around 0.75 even at low cutoffs, indicating that links are found in both directions. In summary, this very simple simulation scenario shows that the use of CI-GWAS marker IVs within an MR framework such as IVW gives robust results, in contrast to selecting the strongest associated markers from a standard GWAS, when comparing at the same p-value threshold.

**Fig. 1:**
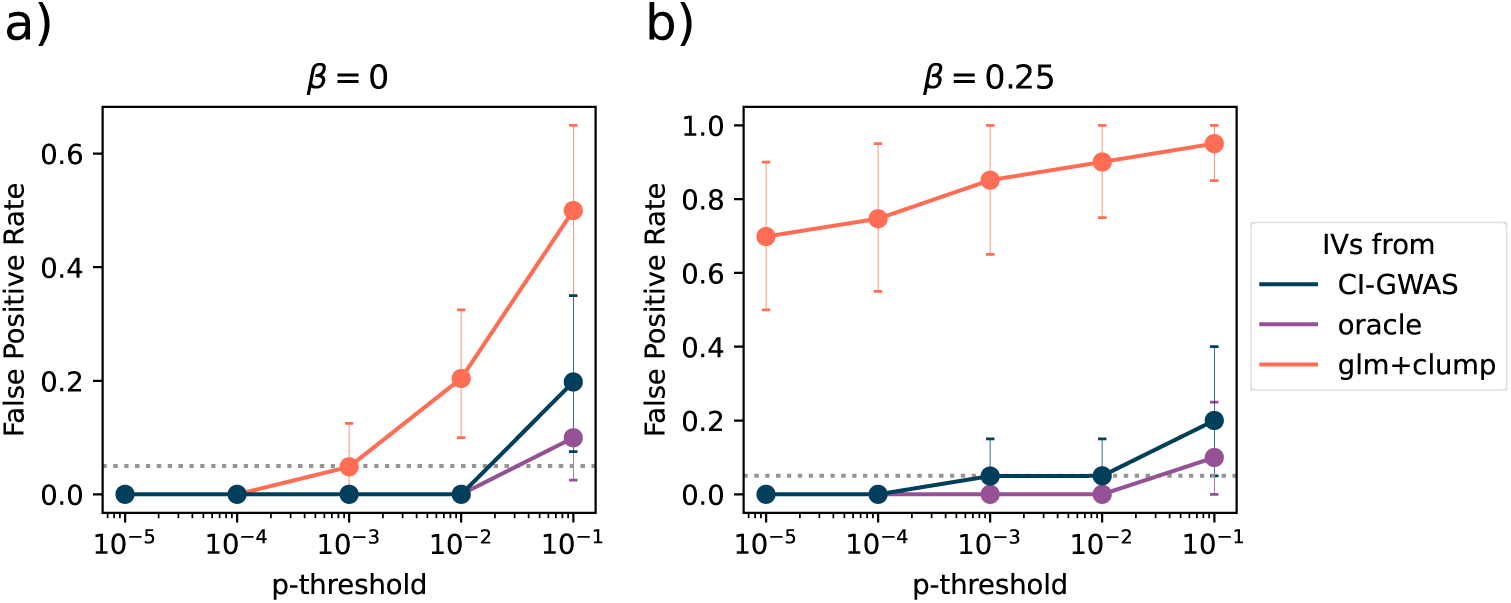
Causal inference performance of IVW with different IV selection in simulation with strong LD and two traits. False positive rate of the inferred causal relationships between the two simulated traits *y*_1_ and *y*_2_, for simulations with a true causal effect *β* of *y*_1_ onto *y*_2_ of (a) *β* = 0 and (b) *β* = 0.25, shown at different p-value thresholds on the x-axis. We compare the performance of IVW supplied with the true markers (oracle), parental markers of traits in the CI-GWAS inferred skeleton or the baseline MR procedure of clumped results of linear models at a p-value threshold for marginal association of 10^−8^ (glm + clump). Gray dotted lines indicate FPR=0.05. Points show bootstrapped means, error bars indicate the bootstrapped 95% confidence interval of the mean (1000 samples), calculated across 20 replicates. A detailed description of all calculations and MR methods used is given in the Methods section.

We then expanded our simulation study to a larger scale by simulating a complex scenario with many correlated traits and latent confounding. We use a structural equation model consisting of twelve continuous traits, two of which are latent confounders and are not used in the inference, each with a set of 50 causal markers chosen at random from 654,173 imputed genetic markers recorded on 458,747 UK Biobank individuals. We simulate marker-trait and trait-trait adjacencies, causal effects, and trait data.

We first confirm the recovery of the randomly drawn underlying causal SNP variants in the presence of real-data LD. We find a low false discovery rate (FDR) of ≈ 0.1 for detecting the exact associated SNP with CI-GWAS at *α* = 10^−3^ rather than selecting an LD proxy SNP (Figure 2a). The true positive rate (TPR) of the associations in the set of selected SNPs is significantly greater for CI-GWAS (≈ 0.6) as compared to selecting the top GWAS variant (≈ 0.4). While this is obvious as GWAS associations do not control for LD, the key comparison is that the number of marker associations inferred per trait are roughly the same for both CI-GWAS and a standard GWAS at standard significance thresholds (Figure S2). This shows that for markers obtained from CI-GWAS, one is more likely to get a fuller set of associations with the traits of interest, compared to simply selecting the top GWAS variant.

**Fig. 2:**
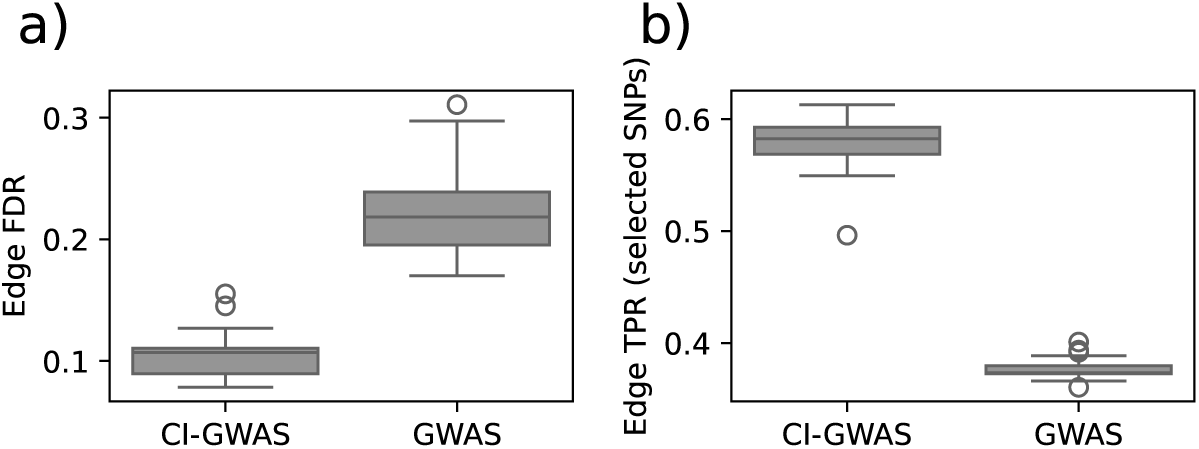
Properties of selected associated variants in CI-GWAS vs GWAS. We compare (a) the false discovery rate (FDR) of SNP-trait associations (edges), as well as (b) the true positive rate (TPR) of the associations in the subset of selected markers between CI-GWAS and GWAS selected SNPs.

In the simulations for this figure, we drew marker-trait effects from *U* [(−10^−2^, −10^−4^) ∪(10^−4^, 10^−2^)], a 0.2 chance for each marker to have a pleiotropic effect on an additional trait (*γ* = 0.2). We run CI-GWAS with an *α* = 10^−3^ and conduct a GWAS with a significance threshold of 10^−8^ for marker selection, followed by clumping to select focal variants. We show data from 20 simulations. Boxes show the interquartile range and the median, whiskers show 1.5× the interquartile range. Datapoints outside that range are shown explicitly.

Next, we evaluate the estimation of the causal effects among traits in comparison to a selection of frequently used MR approaches. We compare three commonly used MR methods, CAUSE [11], MV-PRESSO [8] and MVIVW[9] when used normally and in conjunction with CI-GWAS. To run each of the three MR methods, we use markers selected by standard marginal GWAS testing (MR (*IV_G_*)) or CI-GWAS (MR (*IV_CIG_*)) as input. We then optionally filter the outputs using the CI-GWAS learned skeleton (+ adj) or the PAG, so that we only conduct MR testing for trait pairs that have an adjacency in the skeleton or a corresponding directed edge in the PAG. We apply a significance threshold of *p* ≤ 10^−8^ in the GWAS and a *α* = 10^−3^ for CI-GWAS, which leads to approximately the same numbers of SNPs selected for each trait (Figure S2).

We apply p-value thresholds in the range 10^−8^ to 10^0^ to the MR results to create TPR-TDR curves and also compare approaches at a standard *p* ≤ 0.05 threshold. We show the bootstrapped means across the 20 replicates and 95% confidence intervals of the means in Figure 3.

**Fig. 3:**
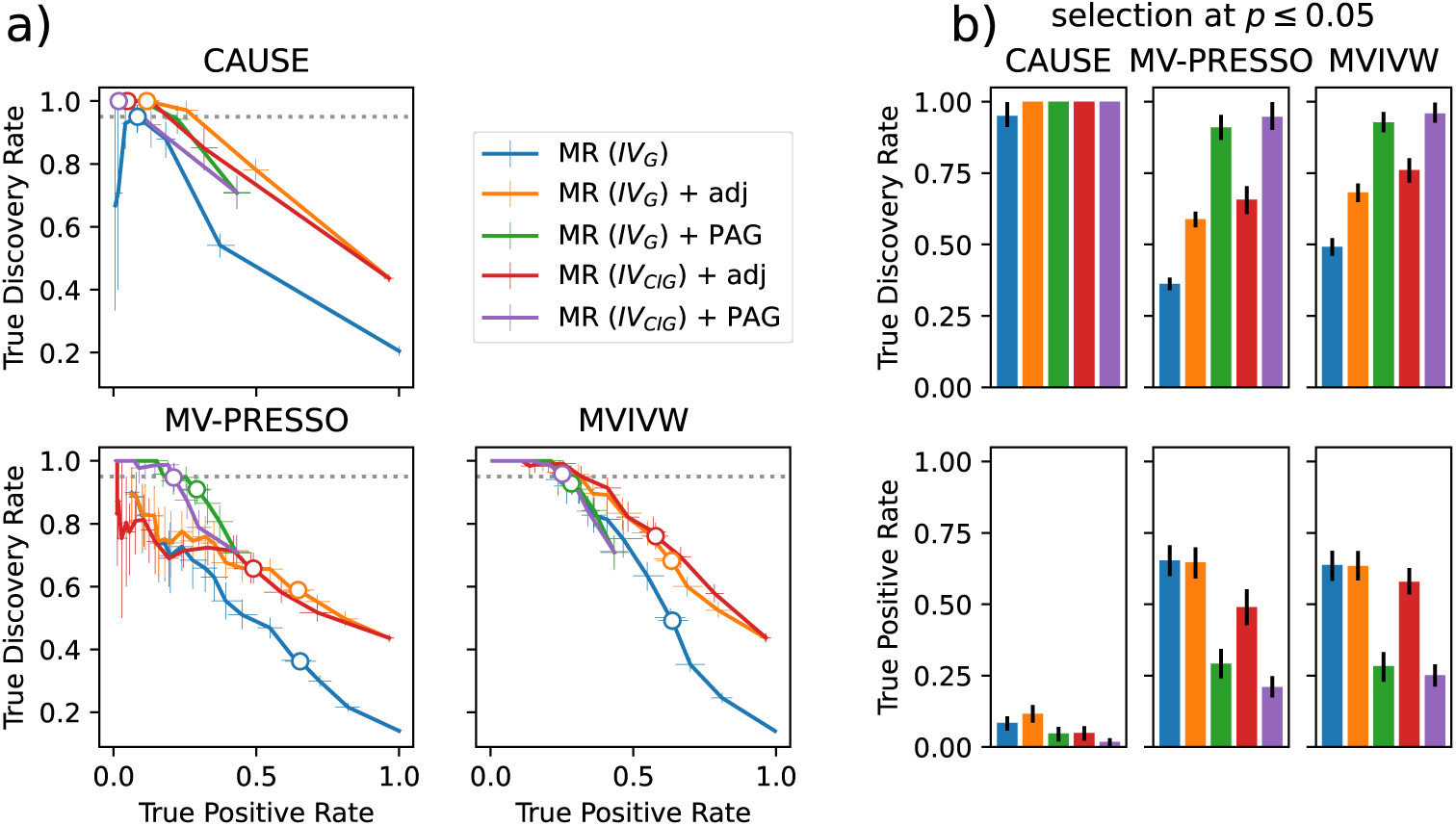
Causal inference performance of CI-GWAS with commonly used MR methods in simulation with strong LD and ten strongly correlated traits. (a) shows the true discovery rate and true positive rate of the trait-trait causal link detection for a number of commonly used MR methods when used with GWAS selected instrumental variables (*IV_G_*), CI-GWAS selected instrumental variables (*IV_CIG_*), and with and without using the CI-GWAS skeleton (adj) or PAG as an additional filter for causal relations. In the simulations for this figure, we drew marker-trait effects from *U* [(−10^−2^, −10^−4^) ∪ (10^−4^, 10^−2^)], a 0.2 chance for each marker to have a pleiotropic effect on an additional trait (*γ* = 0.2). We select *IV_CIG_* and learn the causal graph skeleton and PAG at a threshold *α* = 10^−3^. We select *IV_G_* in a GWAS with a significance threshold of 10^−8^. We used the p-value thresholds in the range [10^−8^, 10^0^] to select significantly causal links from the MR tests. Lines show bootstrapped means, error bars indicate the bootstrapped 95% confidence interval of the mean (1000 samples), calculated across 20 replicates. The dotted line indicates *TDR* = 0.95. Grey filled circles indicate the mean at a p-value threshold of 0.05 and in (b) these values are directly compared across the different modeling choices. A detailed description of all calculations and MR methods used is given in the Methods section.

We make the following observations. First, filtering standard MR results with the CI-GWAS-learned skeleton clearly increases the TDR for all three tested MR methods (MR (*IV_G_*) vs. MR (*IV_G_*) + adj in Figure 3). Second, the TDR is further increased, at the expense of TPR, by using the PAG instead of the skeleton for filtering (MR (*IV_G_*) + adj vs. MR (*IV_G_*) + PAG in Figure 3). Third, for methods such as MV-PRESSO and MVIVW, only by using the PAG for filtering is an FDR of 0.05 achieved at a p-value threshold of 0.05. Fourth, as CAUSE conducts variable selection of the marginal GWAS associations to select IVs, we find no evidence for improved performance when restricting CAUSE to using only CI-GWAS selected IVs. However, inference is more precise, with higher power when using the CI-GWAS-learned skeleton. It should be noted here that CAUSE on its own is not a multivariate MR methods and as such does not distinguish between direct and indirect effects. We take this into account by comparing the CAUSE results to the matrix of true causal paths in the model, and comparing the CAUSE + adj and CAUSE + PAG results to the matrix of true causal direct effects. We would like to emphasize that using CAUSE in conjunction with CI-GWAS offers a way to make the distinction between direct and indirect causal effects. Finally, MVIVW with the PAG learned by CI-GWAS and CI-GWAS selected IVs has a comparable FDR of 0.05 to CAUSE, but has significantly improved TPR. We find that these results remain largely consistent when the selection thresholds are relaxed to *α* = 10^−2^ and a significance threshold in the GWAS of 10^−4^, when more markers are selected (Figures S5 and S4) and when pleiotropic effects are removed from the simulation (Figures S3 and S5). Thus, we show that learning a graphical model, as in CI-GWAS, can be used in conjunction with traditional MR methods to produce methods with improved precision.

### Application to the UK Biobank

Finally, we applied CI-GWAS to 15 traits and 8.4 million single nucleotide polymorphism (SNP) markers in UK Biobank data, as described in the Methods and Table S1. To compare our results to previous findings, we first analyse 36 pairs of risk factors and outcomes (single r.f.), focusing on four outcomes: asthma (AT), cardiovascular disease (CAD), stroke (ST), and type-2 diabetes (T2D). We then analyse all risk factors jointly with each outcome within an MVIVW framework (joint r.f).

The SNP markers discovered in the skeleton of part 1 of our algorithm represent joint marker-trait associations that are conditional on other traits and markers, with the conditioning on other markers controlling for LD (Figure 4a to d). From the single r.f. analyses, we find overall 1052 unique markers, which declines to 696 in the joint r.f. analyses. This suggests that there are few SNPs that are suitable IVs. We also find that the composition of the SNP sets varies between the analyses (Figure S6), as expected due to the varying correlation between the traits considered. Interestingly, we find that the majority of the causal SNPs identified are located in protein-coding genes (Table 1).

**Fig. 4:**
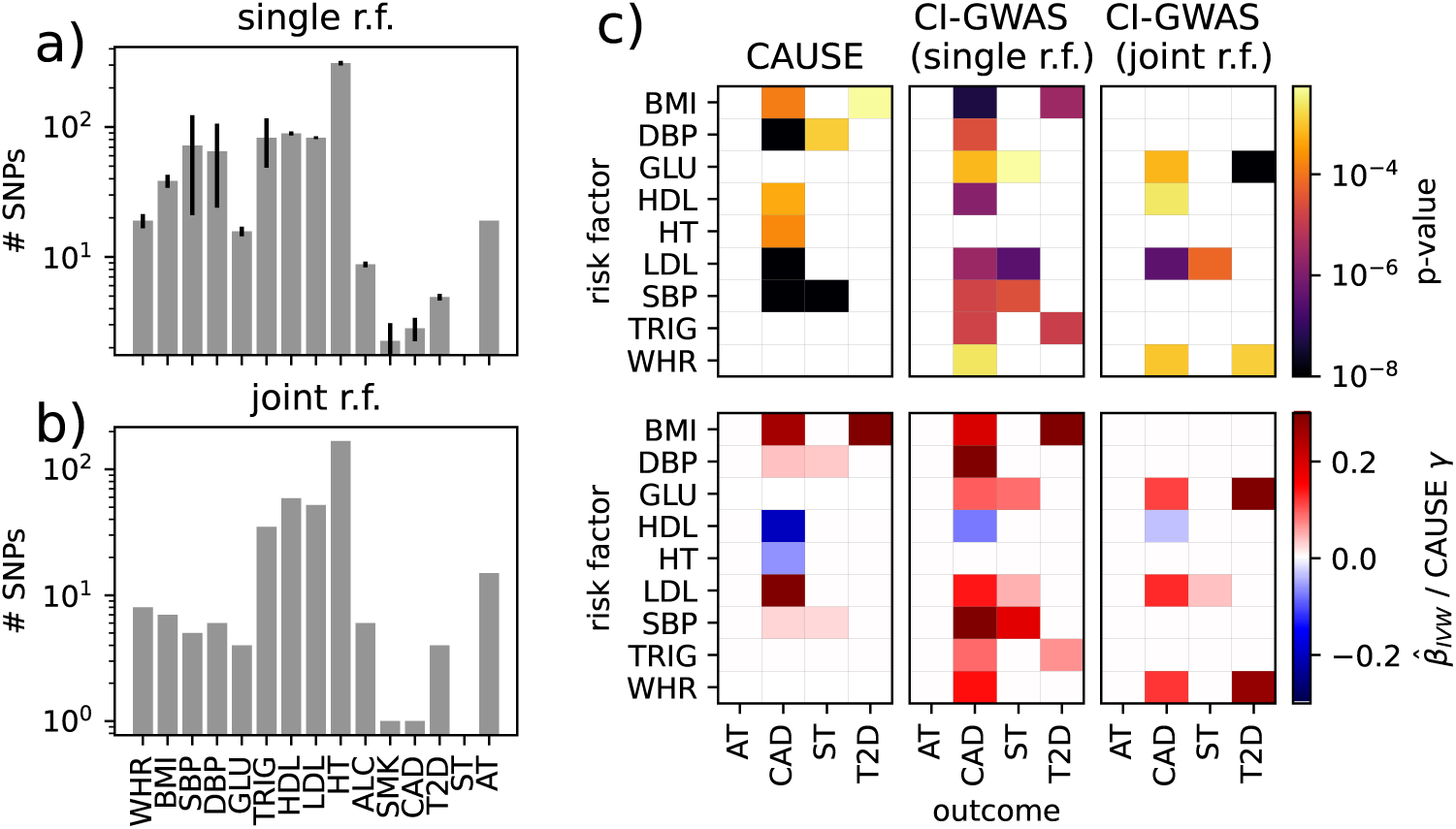
Causal inference in the UK Biobank with CI-GWAS. We analysed 36 risk factor (r.f.) - outcome relationships in the UK Biobank. We ran either pairs of a single risk factor and outcome (single r.f.), or all risk factors with a single outcome jointly (joint r.f.). (a): Numbers of identified causal SNPs per trait in the single r.f. analyses. (b): Number of identified causal SNPs per trait in the joint r.f. analyses. (c): p-values (top row) and causal effect sizes (bottom row) for the analysed risk factor to outcome relations. CAUSE results are taken from [11]. We discard results with p-values above 0.005 (0.05 adjusted for the number of tests) as insignificant, corresponding squares are left white. We use the MVIVW (*IV_CIG_*) + adj approach, selecting only links as causal if the MVIVW result is significant and the traits are adjacent in the inferred skeleton.

**Table 1:** Counts of overlaps of causal SNPs identified by CI-GWAS in the UK Biobank with biotypes of various kinds.

| Biotype | Count |
| --- | --- |
| None | 343 |
| Protein coding gene (exon) | 175 |
| Protein coding gene (intron) | 434 |
| lincRNA (intron) | 53 |
| lincRNA (exon) | 4 |
| Antisense of protein coding gene intron | 36 |
| Antisense of protein coding gene exon | 4 |
| snoRNA | 1 |
| Sense overlapping | 1 |
| miRNA | 1 |

We compare our single r.f. results to those obtained by CAUSE in the UK Biobank presented in [11]. IVW with CI-GWAS skeleton and CI-GWAS identified IVs (CI-GWAS single r.f.) recovers all of the suggested statistically causal paths found by CAUSE, with the exception of height (HT) to CAD and diastolic blood pressure (DBP) to ST. In addition, we discover paths from blood glucose levels to both CAD and ST, from triglycerides to CAD and T2D, and waist-hip ratio (WHR) to CAD (Figure 4e). A causal path is simply a pair-wise relationship between variables, but there remains the potential that the relationship may be explained by other measured/unmeasured variables. Thus, we then test which paths identified in the single r.f. analysis reflect direct adjacencies through a joint r.f. analysis, using MVIVW with CI-GWAS skeleton and CI-GWAS identified IVs (CI-GWAS joint r.f.). Interestingly, CI-GWAS joint r.f. finds GLU and WHR as direct risk factors for both CAD and T2D. HDL and LDL are also direct risk factors for CAD and we find that LDL is also a direct risk factor for ST.

When we filter the exposure-outcome relationships more conservatively using the PAG (as done for MR (*IV_CIG_*) + PAG in Figure 3), we find that only the relationship between GLU and T2D persists (Figure S7). In addition, we find a directed edge from HDL onto AT, for which there is support in neither the previous CAUSE nor in our MVIVW results, but substantial support in the literature. We find that many relationships that are significant in MVIVW are assigned a bidirected edge, which is commonly defined as the absence of a causal relationship and the presence of a latent confounder (Figure S7). Thus, within the UK Biobank, our investigation of potential statistically causal relationships using the PAG cannot rule out the possibility that there is some other unmeasured variable that can explain the relationships we find. This suggests that phenotypic relationships could be predominantly underlain by processes like population structure, linkage, shared parental effects, shared environment and/or assortative mating. Thus, assuming the validity of the assumptions for causal discovery in the UKB data, such as acyclicity of the causal structure, we do not find evidence that blood lipid levels are directly causal for CAD. Rather we find that they may have indirect causal paths to CAD. In contrast, we find evidence that the relationship of blood glucose levels and type-2 diabetes is directly causal.

## Discussion

Here, we present a method for statistical causal inference in a genomic setting to learn a graph by inferring the adjacencies from partial correlations, orientate the edges and then estimate the strength of the causal effects, whilst allowing for the presence of multiple confounders and multiple correlated phenotypes. The algorithms we present have strong theoretical support and have been heavily studied in the causal inference literature [19, 20, 26]. We demonstrate that extending statistical causal discovery from MR methods to multivariable graphical inference for large GWAS or biobank data sets with millions of variables and hundreds of thousands of records provides: *(i)* appropriate selection of IVs that fulfill MR assumptions, and *(ii)* robust inference of potential causal links with greatly reduced false discovery rates and fast run time for both continuous and binary measurements. Current MR methods generally give consistent results, however they all build upon each other, assume a similar fixed directed acyclic graph and can have high type-I error. Here, we demonstrate this empirically and show how this can be addressed by simply combining existing MR approaches with graphical inference.

High-dimensional non-sparse graphical causal inference is an active area of research and there have been a number of proposed alternatives to the approach presented here. Various alternative forms of multivariable MR [8, 27, 28] have been proposed for this problem, but here we show that it is ineffective without being paired with graphical inference. Structural equation models have also been suggested [29], but these require a correct causal order of variables to be known (or learned) and are cumbersome to run on large-scale high dimensional data, even in summary statistic form. A number of artificial intelligence approaches have recently been presented for causal inference in life science settings [30]. However, these lack rigorous theory and an understanding of the conditions required for these algorithms to recover meaningful causal structures and estimates. As a result, while estimates are obtained, one has little assurance that reliable inference is obtained, much like standard MR approaches. Thus, we focus here on applying and developing a principled, theoretically supported method that provides a crucial step toward high-dimensional statistical causal learning for genomics.

A number of limitations remain. Our approach requires either: (i) individual-level data, or (ii) summary information on the minor allele frequency, the LD correlations, and the phenotypic correlations, alongside standard GWAS regression coefficient estimates. This level of summary data are not always publicly available, but could easily be shared, inferred across biobanks, or approximated. Data sets have structure that can inhibit unbiased causal inference such as selection biases like the ascertainment of healthy older people into biobank studies, which can bias causal inference [31]. Additionally, there is also a general tendency in UK Biobank sampled individuals for almost everyone to be under some form of medical intervention. In particular, natural variation in lipid levels and blood pressure may be impossible to observe within the UK Biobank as the majority of people are being treated with some form of drug. Here, we adjust for the effects of drug taking prior to analysis, but if the majority have interventions this is likely ineffective and a larger-scale model investigating prescriptions, observations and outcomes through repeated measures may be more effective. By utilising longitudinal records, where discovered adjacencies in cuda-skeleton can then be orientated longitudinally and combined with therapeutic information, statistical causal effects may then be more effectively examined as there is certainty about the directions of many time ordered edges. This will then reduce the number of bi-directed edges learned in the PAG. Our approach easily facilitates this and we are actively working on this topic. Finally, genotype-phenotype associations likely reflect a complex combination of direct, parental genetic and epigenetic effects, and this may also be why we infer a large number of relationships to occur due to latent confounding. Parental genotypes and their relationship to offspring traits could also be accommodated within our framework but at this time there is a lack of large-scale data.

In summary, our approach is applicable to screen for potential causal relationships within life-science data, from genomics, to the microbiome, to single-cell sequencing and phenotyping datasets. This may be very fruitful as trait-”omics” associations are often stronger than marker-trait correlations, giving improved recall rates. By combining methods of genetic epidemiology and statistical causal inference, within algorithms that require only summary statistic correlations from the data, we expect that our work will facilitate extensive investigation of potential causal hypotheses. We show that inferring causal relationships from simple observational biobank data is limited and our results call for caution when applying current MR methods to biobank data. With any dataset, we feel that it is important to understand which relationships hold under which assumptions, and our approach provides a way to screen associations under explicit checks. In this way, more can be learned about the likely underlying nature of phenotypic and genetic correlations within the human population.

## Methods

### CI-GWAS

CI-GWAS takes either individual or summary-level data, learns conditional dependencies between genetic variants and traits and determines whether conditional dependencies among traits can be attributed to a directed causal relationship. It can then estimate, in conjunction with a Mendelian Randomization method, the magnitude and direction of the potential causal effect. A full analysis with CI-GWAS consists of three computational steps: i) a GPU-accelerated algorithm that conducts genetic variable selection and returns an undirected graph that describes the potential causal relationships among markers and traits (called cuda-skeleton); ii) an approach called sRFCI that directs the skeleton edges and thereby infers causal relationships, allowing for arbitrarily many latent variables; and iii) estimation of the magnitude of the potential causal relationships among the discovered exposures and outcomes with a MR method like multi-variable inverse weighted Mendelian Randomisation (MVIVW) [9]. This final part uses the sets of directly associated markers for each trait as instrument variables, which conform to the assumptions of MVIVW. Additionally, we only report exposure-outcome effect estimates for trait-trait conditional dependencies discovered by either cuda-skeleton or sRFCI that are also significant in MVIVW. We provide detailed information on each step of CI-GWAS in the following sections, along with algorithms for their implementation.

CI-GWAS operates on correlation matrices *M ^G^*, *M ^GT^*, *M ^T^* representing correlations between markers, between markers and traits, and between traits, respectively. As described below, we run the first step of the CI-GWAS (cuda-skeleton) in parallel over LD-blocks, each of which has a distinct *M ^G^* matrix. Thus, we first block the genomic input data with parameter set as shown in Table 2. Then we prepare the correlation matrices as follows:

- *M ^G^*: for each input block *B_i_* we obtain the LD-matrix using PLINK 1.9 with the options --r triangle bin 4 and --allow-no-sex, selecting the rsIDs of the markers in the block with the --from and --to options [32].
- *M ^GT^*: we compute the marker-trait correlations using linear regression models. We first obtain linear regression coefficients *β_ij_* for each marker *i* and trait *j* by running PLINK 2.0 on the bed file with the complete marker data and the individual-level trait-data with --glm omit-ref --variance-sta<u>ndardize opti</u>ons [33]. We then compute the Pearson correlation via *ρ_ij_* = *β_ij_* 2 ∗ *f_i_* ∗ (1 − *f_i_*), where *f_i_* is the minor allele frequency of marker *i*.
- *M ^T^*: Here we simply compute the Pearson correlations between the individual-level data trait columns.

**Table 2:** Algorithm parameters and their values in all applications on real data.

| Algorithm | Parameter | Value |
| --- | --- | --- |
| cuda-skeleton | $l_{max}^{(1)}$ | 3 |
| cuda-skeleton | $l_{max}^{(2)}$ | 14 |
| cuda-skeleton | $\alpha$ | $10^{-3}$ |
| blocking | $s_{max}$ | $10^4$ |
| blocking | $\epsilon$ | $10^2$ |
| blocking | corr-width | $2 * 10^3$ |

Note that any choice of approximations of (i) to (iii) is accepted within our algorithm. For example, recent proposals for summary statistic prediction and heritability models are to “whiten”, “shrink”, or “block” the LD matrix, and to use approximations from other publicly available samples, if the original sample matrices are unavailable [34] can all be accommodated.

In order to accommodate binary traits in our framework, we calculate all correlations with the binary traits placed on the liability scale, as suggested by [25]. For this we use polyserial correlations [35] between continuous variables and binary variables, tetrachoric correlations [36] between binary variables, and Pearson correlations between continuous variables. We provide open source code and scripts to run our full analysis (see Code Availability).

### Part 1: Variable selection and inference of adjacencies

**Algorithm 1.**
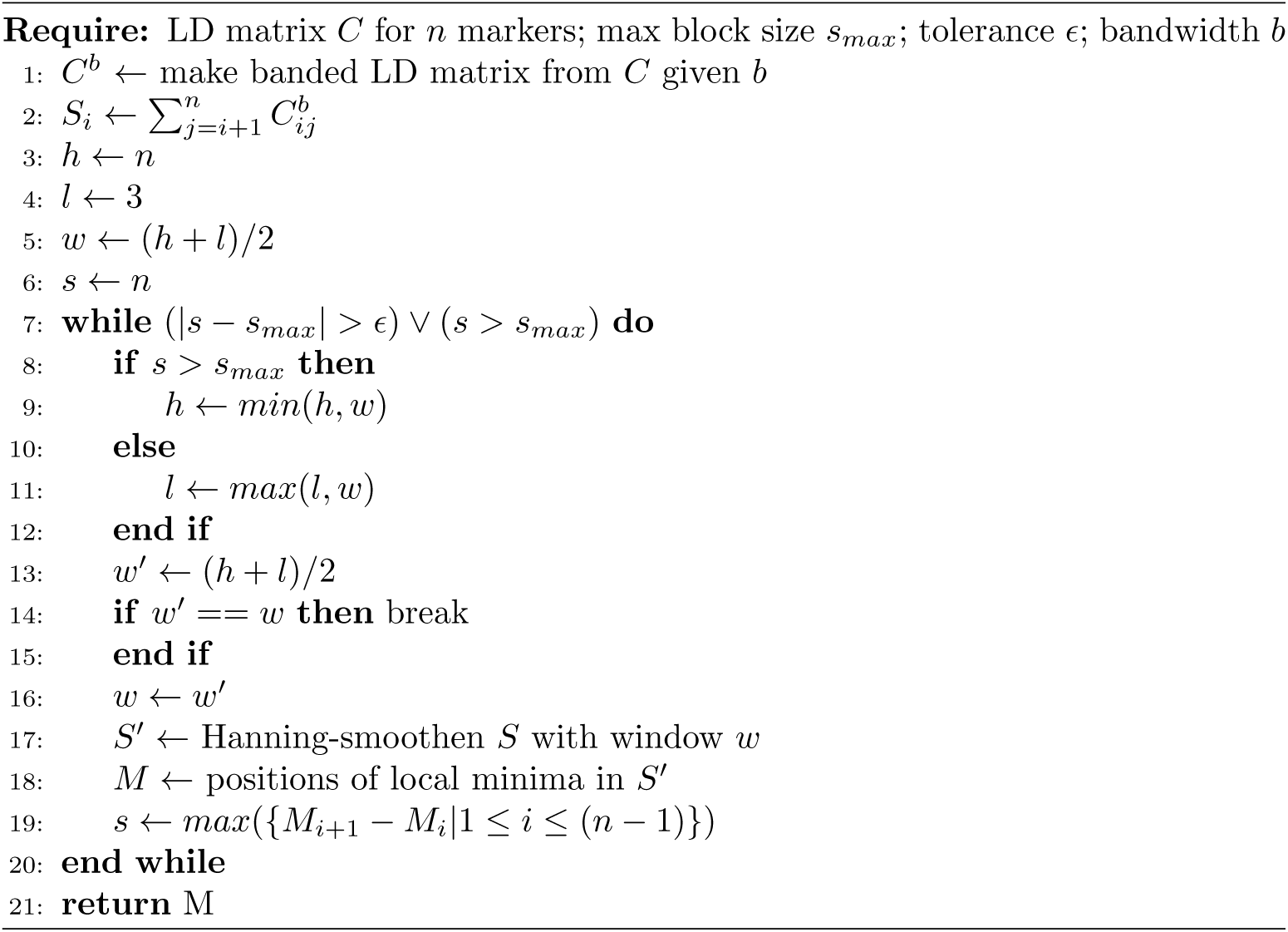
Genome blocking procedure for summary statistics cuda-skeleton

The first step of our algorithm: (i) selects markers that have a statistically significant effect on one or more of the traits; and (ii) builds the *skeleton*: the undirected graph describing all possible direct relationships between all variables (*i.e.* all markers and traits). This step is based on the skeleton inference procedure of the recently proposed cuPC algorithm [21], a GPU-accelerated implementation of the PC-stable algorithm [20].

Given all pairwise correlations between variables, the first part of the PC-stable algorithm infers the skeleton by testing pairs of variables *V_i_* and *V_j_* for independence conditional on an exponential number of *separation sets*, *S*, consisting of other variables in the graph. In particular, *V_i_* ***⊥*** *V_j_*|*S* is tested by first estimating the partial correlation of *V_i_* and *V_j_* given a separation set *S* consisting of neighbors of *V_i_*. We denote this by *ρ*^(*V_i_, V_j_*|*S*). The Fisher’s z-transform of this value, *i.e.*

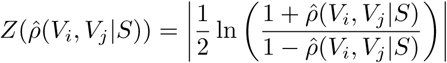

is then compared to a threshold *τ*, given by

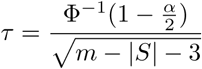

where *m* is the sample size, *α* is the significance level of the test and Φ is the CDF of the standard normal distribution. *Z*(*ρ*^(*V_i_, V_j_*|*S*)) ≤ *τ* is taken to imply *V_i_* ***⊥*** *V_j_*|*S*. These tests are run for consecutively increasing *level l* = |*S*|, starting with *l* = 0. A *l_max_* can be set to limit the runtime of the algorithm. Edges between independent variables are removed. The algorithm terminates when (*l > n*) ∨ (*l > l_max_*), where *n* is the number of variables. The full parameter setting used in this work is given in Table 2.

The runtime scales exponentially with the number of variables, since the number of possible *S* of a given size *l* increases exponentially. cuPC performs the conditional independence tests in a highly parallelized fashion utilizing modern GPU capabilities, enabling skeleton inference for many variables in a short time. In the genomic setting, with order 10^9^ markers and order 10^2^ traits, both storage of the input correlation matrix as well as computation time required, prohibit the application of cuPC on all variables at once. Therefore, we block the marker LD matrix in a fashion inspired by [37] and as shown in Algorithm 1. When relaxing the *α* threshold, there may be circumstances where the memory requirement at larger *l* exceeds that on standard GPU architecture. If this issue arises, we propose to simply run the algorithm twice, first with a low *l* = 3, to select parental markers, and then run all levels from *l* = 0 to *l* = 14 using only the parental markers and all traits. We apply the skeleton inference part of the cuPC algorithm to each block of markers together with all traits individually. We performed a sensitivity analysis for this choice in our empirical UK Biobank application and concluded that this strategy results in approximately the same marker selection that one would obtain from running on full chromosomes, as can be seen in Figure S8.

Having inferred a set of skeletons with adjacency matrices *A^B^* = *A*_1_*, A*_2_*, …, A_n_*and separation sets *S^B^* = *S*_1_*, S*_2_*, …, S_n_*blockwise, we first create a merged skeleton with adjacency matrix *A*^∗^ following four rules: (i) a pair of traits are adjacent in *A*^∗^ if and only if they are adjacent in all *A*^′^ ∈ *A^B^*, since the presence of a single separation is sufficient to imply conditional independence; (ii) if two markers are in different blocks, they are not adjacent in *A*^∗^; (iii) if two markers are in the same block *i*, they are adjacent in *A*^∗^ if and only if they are adjacent in *A_i_*; and (iv) if a marker and a trait are adjacent in any block, they are adjacent in *A*^∗^. This gives a merged skeleton, representing a reduced set of markers that have a direct edge to one or more traits in the graph. This reduces the set of variables to the ones which are most likely to have direct causal effects on traits. In order to prevent fine-mapping errors due to correlated markers being placed in separate blocks, we repeat the cuda-skeleton step on a single artificial block, consisting only of the markers selected in the first run. This ensures that each marker-trait association is always tested conditional on every other potentially correlated marker.

### Part 2: Orientating edges

Having inferred a skeleton, we propose to infer the direction of causal relationships by orientating the graph edges. For this, we employ two procedures: (i) inference of a partial ancestral graph (PAG) by a modified RFCI [19] algorithm; and (ii) multi-variable inverse variance weighted Mendelian randomization (MVIVW) [9], or a comparable method, as described in the following section.

### Inferring a PAG

Unobserved confounding variables (i.e. latent variables) are a common problem for the inference of causal relationships. They indicate violation of causal sufficiency (i.e. knowledge of all causally important variables in the graph), which is a common assumption in causal inference. In consequence, a variable may be inferred as causal for another even though there is no causal relationship between the two. To allow for an additional check for unobserved variables, we devise an approach based on the RFCI algorithm [19] for the inference of causal graphs, which allows for arbitrarily many latent variables in the true underlying directed acyclic graph (DAG). True separation sets, i.e. the full set of variables that render two non-adjacent variables independent when conditioned upon, are difficult to obtain when marker-trait effects are weak. We observed that weak marker-trait effects result in incomplete separation sets, which cause traits to be missing from the separation sets of markers and other traits, which then gives incorrect edge orientation. To solve this issue, we present an order-independent algorithm that uses the RFCI procedure to orientate edges between adjacent variable, whilst considering a locally maximal number of variables in the separation sets used to infer the orientations. Specifically, we infer two types of separation sets: (i) *S^max^*, a maximal separation set, meaning that the addition of any other variable results in *Z > τ*, and (ii) *S^mpc^* a subset of the maximal separation set that locally minimizes the partial correlation between variables *V_i_* and *V_j_*.

*S^max^* is conservative as to what is not included in the separation set, and thus everything that is not within *S^max^* can be considered a collider between the variables. This separation set is used within sRFCI to orientate unshielded triples, as described below. A triple of nodes *V_i_*, *V_k_* and *V_j_* is unshielded, if all of the following are true: (i) *V_i_* and *V_k_* are connected; (ii) *V_k_* and *V_j_* are connected; (iii) *V_i_* and *V_j_* are not connected. We only want to orientate unshielded triples, when *V_k_* is not an element of *S^max^*, and thus is an unlikely collider. The set *S^mpc^* is more conservative with regards to the inclusion of variables into the separation set and we use this in the later orientation steps of sRFCI, as described below.

Algorithm 2 describes the algorithm we created to obtain *S^max^* and *S^mpc^* using the skeleton obtained from cuda-skeleton and a matrix containing the marker-marker correlations, marginal marker-phenotype correlations and phenotype-phenotype correlations. We made *S* order independent and we attempted to overcome missing non-collider variables, by maximising separation sets as much as possible. We note that the CPC/MPC versions of RFCI [20] provide a solution for the same issue by maximally searching all potential separation sets, but this unfortunately is infeasible in a high-dimensional genomic setting as it requires checking 2*^n^* sets where *n* is the degree of *V_i_*. Therefore, we propose the heuristic Algorithm 2, where we do not exhaustively test all possible separation sets, rather we incrementally increase their size, always adding the element that yields the lowest partial correlation among the tested elements at each step. Thereby we reduce the search space to *n*^2^ sets. Furthermore, we reduce the set of unshielded triples that are used for orientation to *relevant unshielded triples*, which we define as those that contain at most one marker, and where the middle variable is not a marker. The separation sets for all other triples are not needed, since we assume that markers temporally precede traits and therefore force marker-trait edges to be directed towards traits. This reduces the number of triples for which separation sets need to be selected greatly, especially when the number of identified markers is large. One limitation of our modifications is that *S^max^* may not be the largest separation set among all possible sets. Likewise, *S^mpc^* may not be the separation set that yields the partial correlation among all possible sets that is closest to zero. However, we found that selecting these separation sets improves the orientation within sRFCI as compared to using the minimal subset *S*^∗^.

We now pass these separation sets to parts of the RFCI algorithms as described below:

Step 1. Create a list of unshielded triples using Algorithm 4.4 of [19].

Step 2. Orientate the v-structures using *S^max^* and Algorithm 4.4 of [19].

Step 3. Make a list of ambiguous unshielded triples *V_i_, V_k_, V_j_*, where the *V_k_* is in *S^max^* but not in *S_i,j_*. These ambiguous triples do not then contribute to the orientation in later stages, although they themselves can later be orientated if there is evidence for this.

**Algorithm 2.**
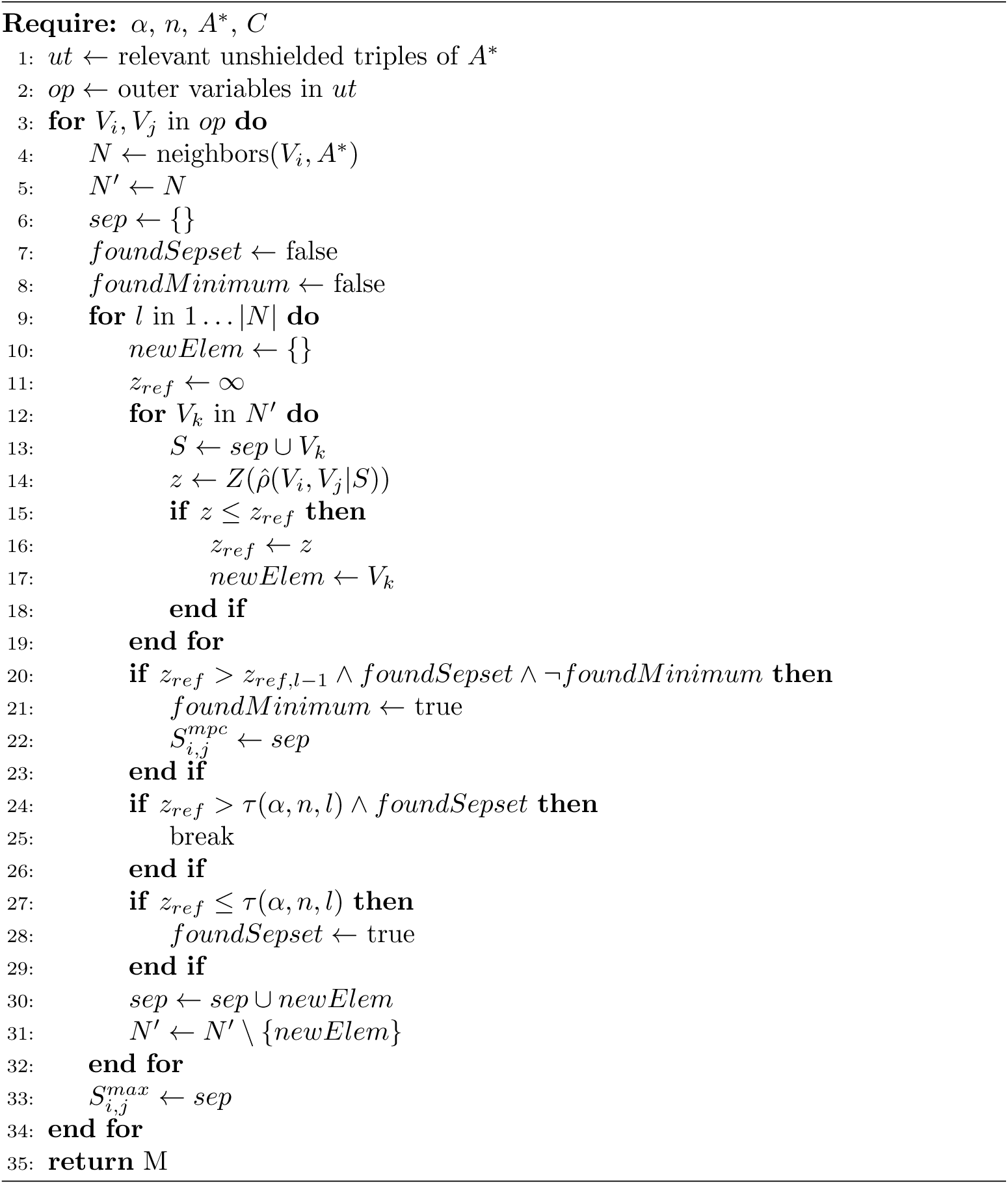
Separation set selection

Step 4. Further edges are oriented as much as possible following the guidelines provided in Algorithm 4.5 of [19]. This step involves repeated applications of orientation rules (R1) to (R10) of [38]. We implemented the suggested modifications from [20] to rules R1-R10 that make them order-independent. In this step, we provide the option to limit the application to only the phenotypic edges in order to reduce the runtime.

Step 5. All marker-phenotype edges are orientated to ensure that phenotypes are not ancestral to markers. This conforms to the assumption that marker states temporally precede phenotypes.

The output of this algorithm is a Partial Ancestral Graph (PAG) representing a the set of possible Maximal Ancestral Graphs (MAG, see Ref. [19] for definition). Adjacencies are inferred to be composed of arrows (*<* or *>*), circles (◦), and tails (−).

An arrow at a variable implies that the variable is not ancestral to the variable at the other end of the edge. The tail implies that the variable is ancestral to the variable at the other end of the edge. Circles represent arrowheads in some MAGs in the Markov equivalence class represented by the PAG and tails in others. RFCI allows inference of causal information in the presence of latent variables and hidden factors or confounders [19], which has been shown to be of importance for Mendelian Randomisation methods applied to biobank data [39, 6, 40, 41, 11]. A situation can arise when neither variable is ancestral to the other, which results in the adjacency being inferred as ←→. This implies that neither of the two variables is ancestral to the other, with no direct causal relationship inferred, implying the existence of a latent confounder.

To select a set of relationships with clear causal direction from the output of this procedure, we select all inferred edges with one arrow and one tail, since these are invariant across the MAGs represented by the PAG and therefore are inferred to have a definite orientation.

### Part 3: Learning the magnitude of the causal effects

Having learned the conditional dependencies in the data with cuda-skeleton and tested for the presence of latent confounding among traits with sRFCI we then propose to use the framework of MVIVW, or a comparable method, to assess the significance and magnitude of causal effects.

In order to identify potential causal relationships in an MR framework like MVIVW, sets of IVs and exposures are needed for each outcome. Specifically, MVIVW assumes that IVs are associated with at least one exposure, are not associated with any latent confounders of the exposures and outcome, and are conditionally independent of the outcome given the exposures, *i.e.* that all causal paths from the IV to the outcome go through the exposures, and that there is no latent confounder of IV and outcome [42, 9]. MVIVW can be employed using the SNPs inferred from cuda-skeleton because the conditional-independence tests conducted in *cuda-skeleton* explicitly check for several common MR assumptions on IVs (see [42] for a discussion of those assumptions). In consequence, valid IVs can be directly read off the skeleton. In particular, for a given exposure trait *A* and an outcome trait *B*, any marker *X* adjacent to *A* that is not adjacent to *B* is a valid instrument to test the effect of *A* on *B* (assuming the skeleton is faithful to the underlying distribution), since

1. if *X* had a direct effect on *B*, the dependency should not have been removed by any conducted test. Thus the exclusion restriction holds.
2. if marker *X* and the outcome *B* relationship was confounded by a latent variable, that confounding would induce a dependency, and again, *X* and *B* should remain adjacent in the skeleton, since no conducted test should be able to remove the dependency. Thus the effect of *X* on *B* should not be confounded.

Note that the above only holds as long as all traits in the model except for the outcome are used as exposures in a multivariate MR analysis. This is necessary because *X* could also be associated with *B* through the omitted trait, which could either have in turn an effect on *B* and thereby invalidate the exclusion restriction. Principled instrumental selection of this kind stands in contrast to most other current MR approaches, where IVs are selected by taking variants with the highest marginal association test statistics. A similar approach, based on the association of markers with a single trait, while having multiple traits in the model, has been proven to be valid and successfully applied in previous work [16]. Here, we advance this idea in a framework where no markers need to be pre-selected, many variables can be conditioned on at once, and pleiotropic markers can be used for inference, promising more accurate IV selection and powerful analysis.

For each outcome, we assess the significance and estimate the effect size of each potential exposure using MVIVW with unfiltered parental markers (conforming to the criteria above) as instrumental variables. We use all traits in the model, aside from the outcome, as exposures. From here, we select only the links that i) are selected as significant in cuda-skeleton or ii) are directed in the same way as inferred by MVIVW in the sRFCI inferred PAG. This gives sets of potential causal links with evidence for a direction that have been learned from the data.

We use the selected instrumental variable and exposure sets as input to the MVIVW implementation in the *MendelianRandomization* R package [43], with the robust option. We select significant causal relationships by applying a p-value threshold of 0.05 to the output.

### Application of CI-GWAS to the UK Biobank data

UK Biobank has approval from the North-West Multicenter Research Ethics Committee (MREC) to obtain and disseminate data and samples from the participants (https://www.ukbiobank.ac.uk/ethics/). These ethical regulations cover the work in this study. Written informed consent was obtained from all participants. From the measurements, tests, and electronic health record data available in the UK Biobank data [44], we selected six blood based biomarkers, four of the most common heritable complex diseases, and seven quantitative measures. The full list of the 17 traits, the UK Biobank coding of the data used, and the covariates adjusted for are given in Table S1. For the quantitative measures and blood-based biomarkers, we adjusted the values by the covariates, removed any individuals with a phenotype greater or less than 7 SD from the mean (assuming these are measurement errors), and standardized the values to have mean 0 and variance 1.

For the common complex diseases, we determined disease status using a combination of information available. For asthma (AT), we used self-report asthma diagnosed by a doctor (3786-0.0), date of asthma report (42014-0.0), and self-report recent medication for asthma (22167-0.0). For cardiovascular disease (CAD), we used self-report information of whether a heart attack was diagnosed by a doctor (3894-0.0), the age angina was diagnosed (3627-0.0), whether the individual reported a heart problem diagnosed by a doctor (6150-0.0), and the date of myocardial infarction (42000-0.0). For stroke (ST), we used self-report stroke diagnosed by doctor (6150-0.0), the age stroke was diagnosed (4056-0.0). For type-2 diabetes (T2D), we used self-report information of whether diabetes was diagnosed by a doctor (2443-0.0), the age diabetes was diagnosed (2976-0.0), and whether the individual reported taking diabetes medication (6153-0.0, 6177-0.0). For each disease, we then combined this with primary death ICD10 codes (40001-0.0), causes of operative procedures (41201-0.0), and the main (41202-0.0), secondary (41204-0.0) and inpatient ICD10 codes (41270-0.0). For AT, we selected all ICD10 J45 codes, for CAD we selected ICD10 codes I20-I29, for ST we selected all ICD10 I6 codes, and for T2D we selected ICD10 codes E11 to E14 and excluded from the analysis individuals with E10 (type-1 diabetes). Thus, for the purposes of this analysis, we define these diseases broadly simply to maximise the number of cases available for analysis. Individuals with neither a self-report indication or a relevant ICD10 diagnosis, were then assigned a zero value as a control.

We first restricted our analysis to a sample of European-ancestry UK Biobank individuals. To infer ancestry, we used both self-reported ethnic background (UK Biobank field 21000-0), selecting coding 1, and genetic ethnicity (UK Biobank field 22006-0), selecting coding 1. We projected the 488,377 genotyped participants onto the first two genotypic principal components (PC) calculated from 2,504 individuals of the 1,000 Genomes project. Using the obtained PC loadings, we then assigned each participant to the closest 1,000 Genomes project population, selecting individuals with PC1 projection ≤ absolute value 4 and PC2 projection ≤ absolute value 3. We applied this ancestry restriction as we wished to provide the first application of our approach, and to replicate our results, within a sample that was as genetically homogeneous as possible. Note however, that our approach can be applied within different human groups (by age, genetic sex, ethnicity, etc.). Our future work will focus on exploring differences in causal inference across a diverse range of human populations.

Samples were also excluded based on UK Biobank quality control procedures with individuals removed of (i) extreme heterozygosity and missing genotype outliers; (ii) a genetically inferred gender that did not match the self-reported gender; (iii) putative sex chromosome aneuploidy; (iv) exclusion from kinship inference; (v) withdrawn consent. We used genotype probabilities from version 3 of the imputed autosomal genotype data provided by the UK Biobank to hard-call the single nucleotide polymorphism (SNP) genotypes for variants with an imputation quality score above 0.3. The hard-call-threshold was 0.1, setting the genotypes with probability ≤ 0.9 as missing.

From the good quality markers (with missingness less than 5% and p-value for Hardy-Weinberg test larger than 10^−6^, as determined in the set of unrelated Europeans), we selected those with minor allele frequency (MAF) ≥ 0.0002 and rs identifier, in the set of European-ancestry participants. This resulted in data set with 458,747 individuals and 8,430,446 markers. We apply CI-GWAS to these data as described above and we use these data for our simulation study as described below.

### Simulation study

We provide code to simulate data of the kind presented in either of the simulation scenarios (see Code Availability).

### Simple scenario

We set up a structural equation model (SEM) of two continuous trait variables *A* and *B* with 200 causal markers *X_A_*, *X_B_* each, chosen randomly from the 654,172 markers on chromosome 1 in the 8,430,446 marker UK Biobank data set. To model pleiotropy, we allow allow an intersection of *X_A_* and *X_B_*of size 200 × *γ*, where *γ* ∈ {0.0, 0.5}. We simulate two scenarios: one where there is no causal effect between traits *A* and *B* (*β* = 0) and one where *β* = 0.25. The effects of the markers in *X_A_*, *X_B_* on the traits are sampled such that the heritability of each trait is equal to 0.2. We repeat this SEM generation for 20 distinct replicates for each combination of values of *γ, β*. Given a sampled SEM and the marker data from UKB we then sample values for traits *A* and *B* for 458,747 individuals.

We compare the TDR of the causal relationships among the two traits between MVIVW [9] (from the MendelianRandomization R package (version 0.4.2) [45]) when run with GWAS selected markers (as described above), CI-GWAS selected markers (as described above) or oracle markers, *i.e.* the true causal markers from the SEM. We apply range of p-value thresholds (10^−5^, 10^−4^, 10^−3^, 10^−2^, 10^−1^) to select the positive test outcomes from MVIVW. For all metrics calculated for this simulation we present the bootstrapped mean and its 95% confidence interval, calculated in 1000 draws with replacement from the 20 replicates.

### Complex scenario

To simulate a more complex setting, we follow a similar strategy as described above for the simple setting. For twelve continuous traits, we select 50 causal markers each using imputed SNP data from chromosome 1 from the same data as described above. We then simulated a marker-trait and trait-trait adjacency matrix *G*, causal effects *A*, and phenotypic data as described below.

First, we simulate a randomly directed acyclic graph G and generate synthetic data based on the specified graph structure, with the following parameters: the number of observations in the dataset (*n*), the number of SNP variables (*m*), the number of treatment variables (*p*), the number of latent variables (*l*), the average node degree (*deg*), the probability of pleiotropy (a SNP affecting multiple traits) (*probPleio*), and bounds for the causal effect sizes (*lo* and *hi*). The following steps are used to generate the data:

Step 1. Initialize a graph structure represented by a sparse adjacency matrix **G** of dimensionality *d* × *d*, where *d* = *m* + *l* + *p*. Set all elements of **G** to zero. Here we chose *l* = 2; *p* = 10.

Step 2. Generate random adjacencies by setting values of **G** to 1 with probability *<u>^deg^</u>*.

Here we chose average degree *deg* = 3.

Step 3. Create additional marker-trait links, by iterating over each SNP variable and randomly connecting it to latent and phenotype variables with a probability of pleiotropy (*probPleio*). The connections are added to the adjacency matrix **G** accordingly. Here, we selected *probPleio* = 0.2 or 0.0 for 20 replicates each.

Step 4. Given the established adjacency matrix, sample the causal effects for the edges from a uniform distribution within specified bounds (*lo* and *hi*), and store them in matrix **A** as a true causal effect matrix. Here, we used a uniform distribution *U* [(−0.01, −0.0001) ∪ (0.0001, 0.01)] to set the causal effect sizes between markers and traits, and a uniform distribution *U* [(−0.2, −0.001)∪(0.001, 0.2)] between traits.

This approach allowed us to simulate a wide range of causal effects, and to have separate strengths for marker-trait and trait-trait relationships, which reflected the pattern we observed in the UK Biobank data.

Step 5. Simulate data based on a structural equation model. For each variable *V_j_* in **G**, check if *V_j_* has parents (*Pa*(*V_j_*)) in **G**. Consider only variables with an index *i < j* as parents. If *V_j_* has no parents, assign it random values drawn from a standard Normal distribution. If *V_j_* has parents, a linear regression model is constructed using the parental variables’ values. In other words, apply the following structural equation model to generate *V_j_*:

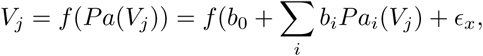

where *b*_0_ is the intercept (set to zero) and coefficients for the regression model are the appropriate *b_i_* values selected from **A** for the *i^th^* parent of *V_j_*. The sampling error *ɛ_x_*is then sampled from a standard normal with variance 1 − *var*( *_i_ b_i_Pa_i_*(*V_j_*)). This results in all variables having unit variance and zero mean, up to a sampling error.

We simulated data for all 458,747 UK Biobank individuals and conducted 40 replicates (20 for each pleiotropy regime), which we analysed using CI-GWAS. For the MR analysis comparison, we used three commonly employed methods: IVW, MR-PRESSO, and CAUSE and their multivariable extensions where appropriate. IVW were implemented using the MendelianRandomization R package (version 0.4.2) [45]. MR-PRESSO was implemented using code from the original publication (version 1.0) [8] with outlier correction. We employed the CAUSE R package (version 1.2.0.0331) [11] to perform the CAUSE analysis. Unless stated we used default options given by each software.

When comparing against MR methods, we supply the methods where appropriate with IVs selected by the standard procedure of running marginal association tests. To perform the marker selection, we run PLINK 2.0 [33] with --glm omit-ref --variance-standardize on the marker data and the trait data. Secondly, we clump the results by running PLINK 2.0 with the options --clump --clump-field P --clump-p1 10e-8 (or --clump-p1 10e-4) and --clump-r2 0.1 --clump-snp-field ID on the results. We then select the focal markers across all clumps for downstream analysis.

To compare the causal inference performances, we calculate TPR and TDR of the returned trait-trait causal relationships, To calculate TPR and TDR, we compare to the directed adjacency matrix of the true DAG. We calculate the sum of all true edges P, which is the total number of possible true edges. We compute TP, by logical AND between the true matrix and the inferred causal relationship matrix. We compute FP, which is the number of inferred, but not true edges. We then calculate TPR = TP/P and TDR = (TP / (TP + FP)). For the TDR calculation for single variable MR, the true matrix differs: it is the matrix of all causal paths in the true graph over the traits. We calculate this because MR estimates causal effects between variables, without regard (a distinction) between edges and paths, or in other words between direct or mediated (through other variables) causality. Multivariable MR (MV-MR) differs as it seeks to infer direct effects and thus for MV-MR we calculate TP, FP, TPR and TDR in the same way as for CI-GWAS.

Since CAUSE estimates path effects instead of direct effects, we compare the estimated effect to the total effect in the true DAG over all path lengths when assessing its performance. The matrix of total effects *M_tot_* is calculated as *M_tot_* = L_n_ *M ^l^*, where *M* is the matrix of the true direct causal effects of the DAG and *n* is the number of traits in the DAG.

### Assessment of the error rate when including binary outcomes in the analysis

We also conducted a simulation to illustrate that treating binary outcomes as continuous can lead to increased error rates when conducting conditional testing of the sort that is done in the PC-algorithm (Figure S1). We conducted two small simulations: (i) independently drawn samples of *x* ∼ *Binom*(2, 2 × 10^−3^) and *y* ∼ *Binom*(1, 0.028). We then tested *CI*(*x, y*) as described above with *α* = 0.05. (ii) we simulated from the DAG: *x* → *y_cont_* → *y_bin_* where *x* ∼ *Binom*(2, 2 × 10^−3^), *y_cont_* ∼ N (*βx,* 1), ∼ N (*γy_cont_,* 1), and *y_bin_* is 1 wherever *y^l^ >* Φ^−1^(*K*) and 0 otherwise. We used *β* = (2 × *MAF* ×(1 − *MAF*))^−0.03^ and *γ* = 0.4. We drew *n* = 458746 samples for each variables, in 10,000 replicates for each scenario. We calculated correlations between the variables in three different ways: “pearson” - Pearson correlations between all variables; “polyserial”: polyserial between (*x*, *y_bin_*) and (*y_cont_*, *y_bin_*); “polychoric”: polychoric between (*x*, *y_bin_*), polyserial between (*y_cont_*, *x*) and (*y_cont_*, *y_bin_*). We calculated the correlations using the ‘hetcor’ function in the ‘polycor’ R package [46].

### Postprocessing of CI-GWAS selected parental markers

Regarding the markers identified as parental to traits in the UKB analysis, we were interested in their potential biological significance, as well as their concordance with previous findings or their novelty.

To address the former, we downloaded ‘Homo sapiens.GRCh37.87.chr.gff3‘ from Ensembl [47], release 110. We then extracted all genes (except for pseudogenes) and their exon positions and sorted the file using

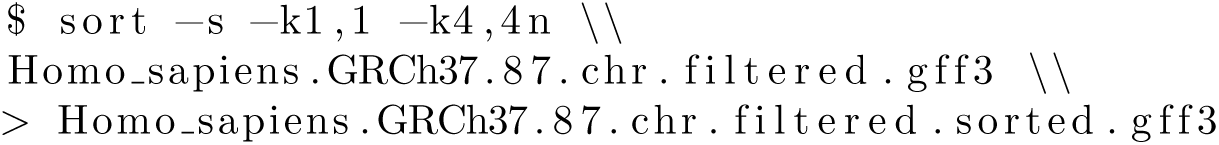

We then searched for the nearest feature to each identified marker using BEDtools [48] version 2.30.0 with

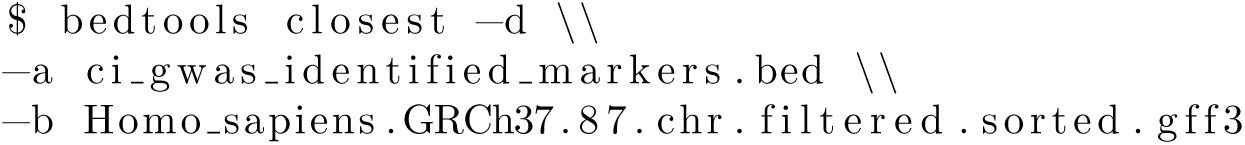

## Ethical approval declaration

This project uses UK Biobank data under project 35520. UK Biobank genotypic and phenotypic data is available through a formal request at (http://www.ukbiobank.ac.uk). The UK Biobank has ethics approval from the North West Multi-centre Research Ethics Committee (MREC). Methods were carried out in accordance with the relevant guidelines and regulations, with informed consent obtained from all participants.

## Supplementary information

Supplementary Information accompanies this work.

## Acknowledgments

We thank Zoltan Kutalik and members of the Robinson group at ISTA for their comments, which improved this manuscript. This work was funded by a research collaboration agreement between Boehringer Ingelheim and the research group of MRR at the Institute of Science and Technology Austria. Additional funding was also provided by an SNSF Eccellenza Grant to MRR (PCEGP3-181181), and by core funding from the Institute of Science and Technology Austria. We would like to acknowledge the participants and investigators of the UK Biobank study. High-performance computing was supported by the Scientific Service Units (SSU) of IST Austria through resources provided by Scientific Computing (SciComp).

## Declarations

### Author contributions

MRR, MM and NM conceived and designed the study, conducted the analysis and wrote the paper. MJB and MRR provided study oversight. IK contributed to study design and analysis. MB contributed to the analyses. All authors approved the final manuscript prior to submission.

### Data availability

This project uses the UK Biobank data under project number 35520. UK Biobank genotypic and phenotypic data is available through a formal request at (http://www.ukbiobank.ac.uk). All summary statistic estimates are released publicly on Dryad: https://doi.org/xx.xxxx/dryad.xxxxxxxxx.

### Code availability

The CI-GWAS code is fully open source and available at https://github.com/medical-genomics-group/ci-gwas. The scripts used to execute the model are available at https://github.com/medical-genomics-group/ci-gwas. R version 4.2.1 is available at https://www.r-project.org/. Plink version 1.9 is available at https://www.cog-genomics.org/plink/1.9/.

## Supplementary Information

**Fig. S1:**
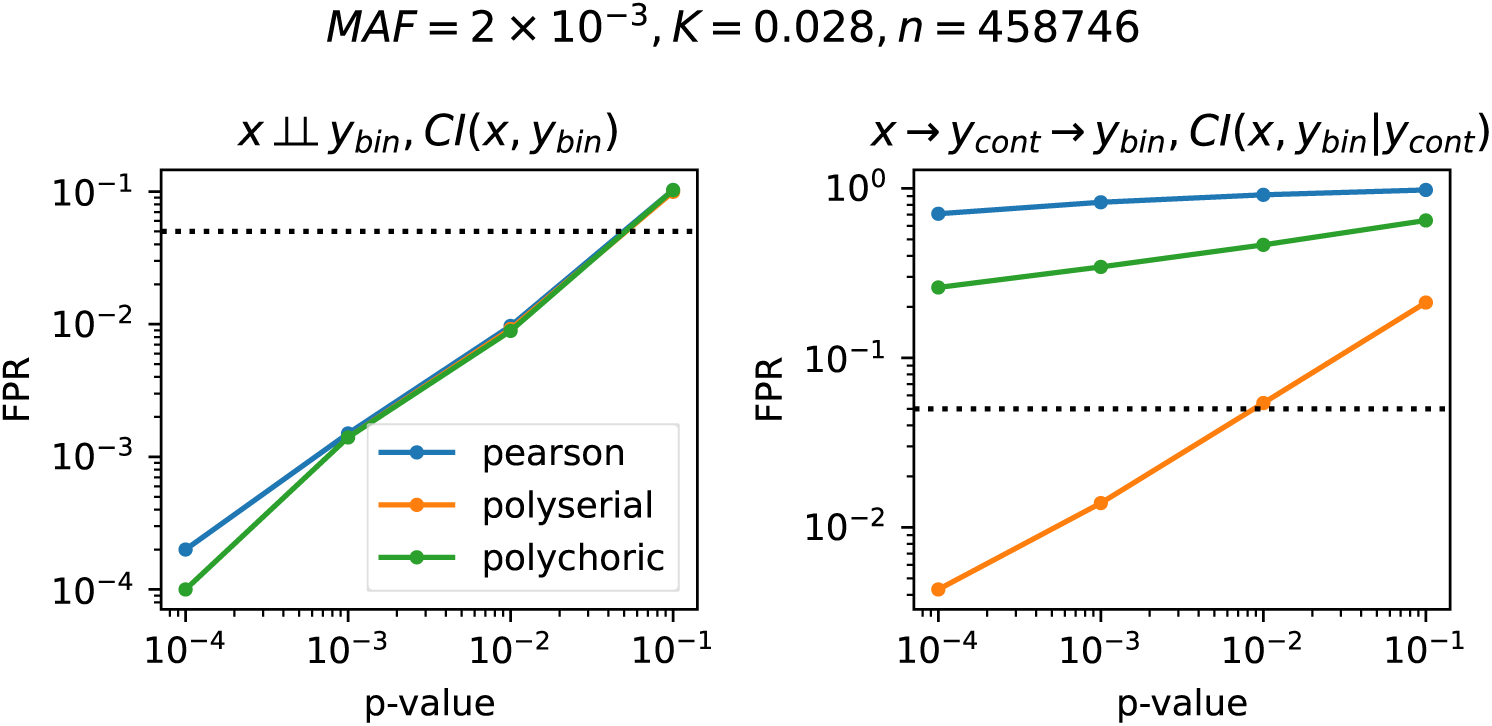
False positive rates when conducting independence tests on ordinal traits based on partial correlations. On the left: independently drawn *x* ∼ *Binom*(2, 2 × 10^−3^) and *y* ∼ *Binom*(1*, K*). On the right: *x* → *y_cont_* → *y_bin_* where *x* ∼ *Binom*(2, 2 × 10^−3^), *y_cont_* ∼ N (*βx,* 1), *y^l^* ∼ N (*γy_cont_,* 1), and *y_bin_* is 1 wherever *>* Φ^−1^(*K*) and 0 otherwise. Here we used *β* = (2 × *MAF* × (1 − *MAF*))^−0.03^ and *γ* = 0.4. We drew *n* = 458746 samples for each variables, in 10,000 replicates for each scenario. We calculated correlations between the variables in three different ways: “pearson” - Pearson correlations between all variables; “polyserial”: polyserial between (*x*, *y_bin_*) and (*y_cont_*, *y_bin_*); “polychoric”: polychoric between (*x*, *y_bin_*), polyserial between (*y_cont_*, *x*) and (*y_cont_*, *y_bin_*). We tested for conditional independence using partial correlations as described in the methods.

**Fig. S2:**
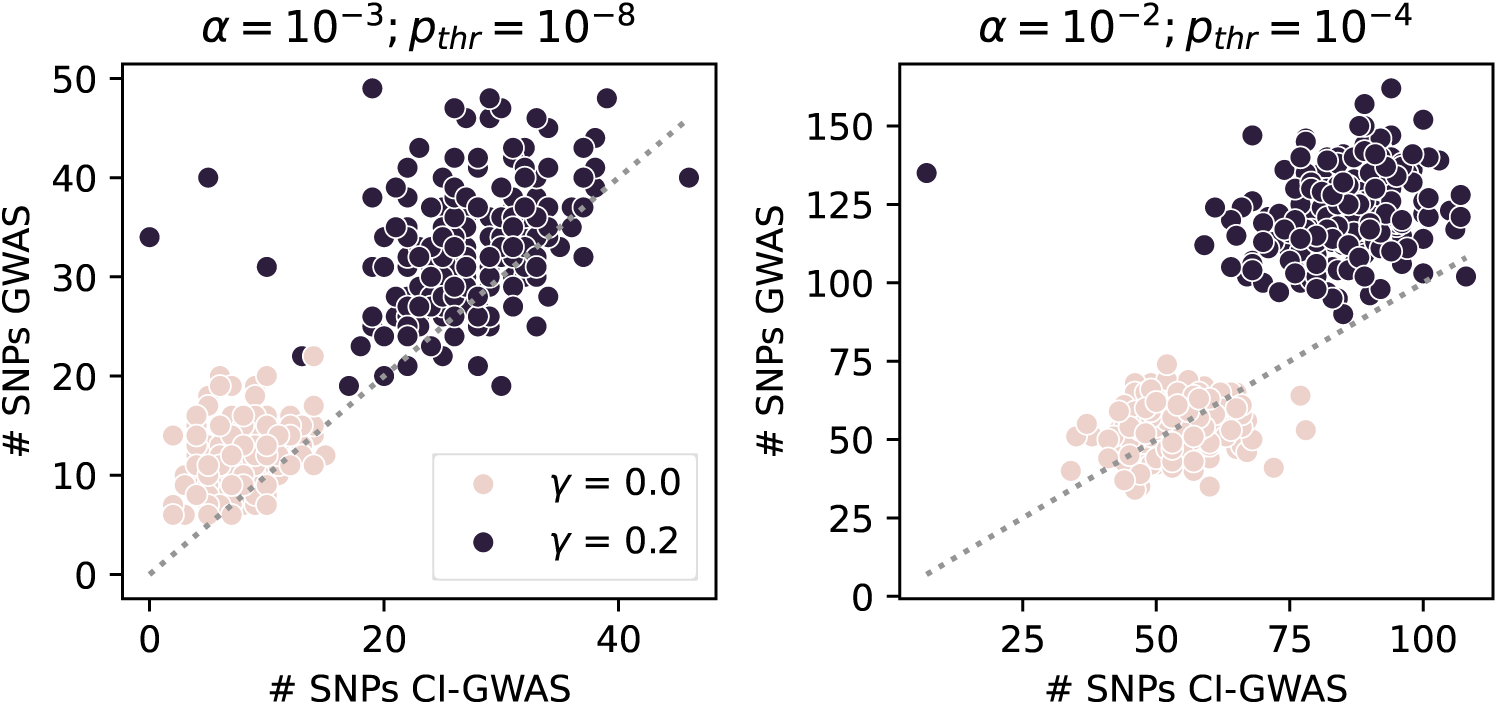
Numbers of variables selected in a GWAS vs CI-GWAS at different significance thresholds. We show the numbers of markers selected for each trait in a simulation with 10 observed and two latent traits in 20 replicates. We simulate two pleiotropy scenarios: no pleiotropy (*γ* = 0) and common pleiotropy (*γ* = 0.2), where *γ* is the chance of each marker to have a pleiotropic effect on a second trait. *α* is the significance threshold in CI-GWAS tests. *p_thr_* is the p-value threshold for selecting markers in the GWAS. The dotted line indicates the diagonal.

**Fig. S3:**
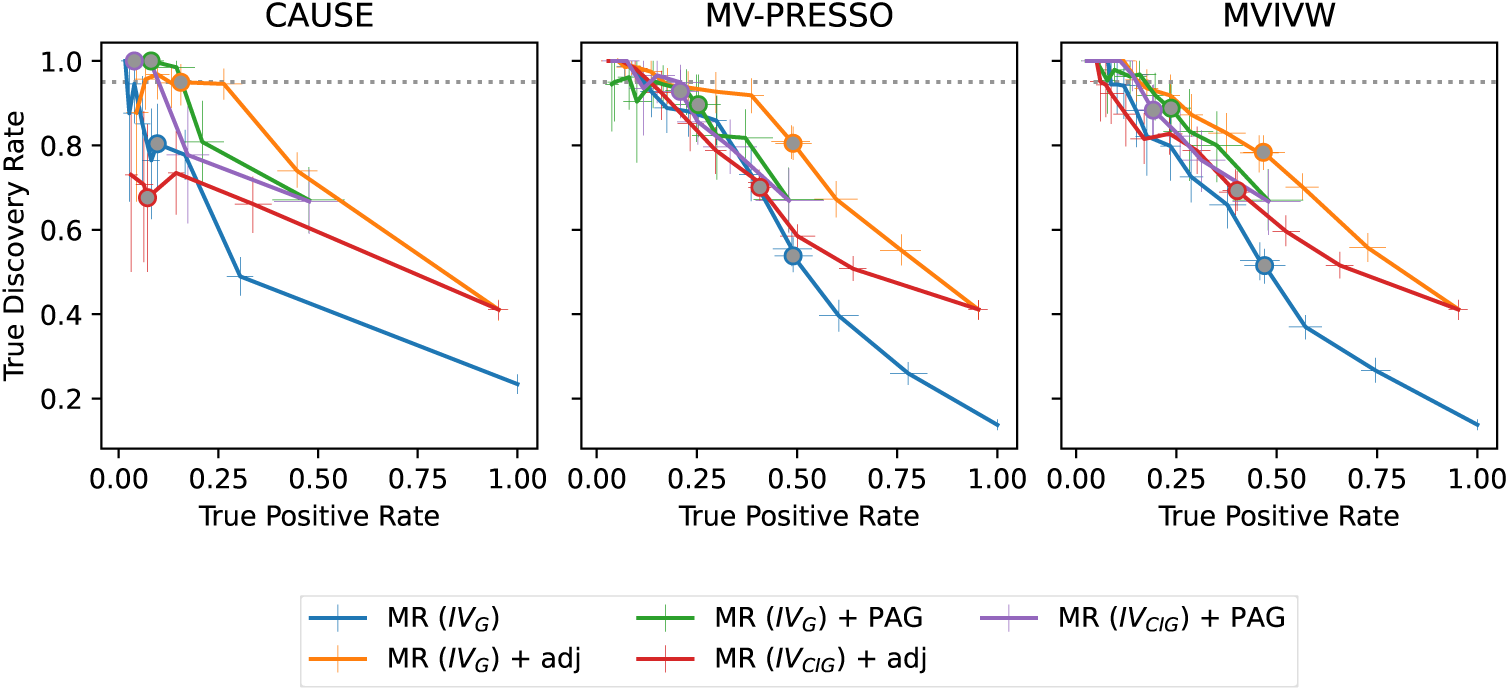
Causal inference performance of CI-GWAS with commonly used MR methods in simulation with strong LD and ten strongly correlated traits without pleiotropy. Here we show the true discovery rate and true positive rate of the trait-trait causal link detection for a number of commonly used MR methods when used with GWAS selected instrumental variables (*IV_G_*), CI-GWAS selected instrumental variables (*IV_CIG_*), and with and without using the CI-GWAS skeleton (adj) or PAG as an additional filter for causal relations. In the simulations for this figure, we drew marker-trait effects from *U* [(−10^−2^, −10^−4^) ∪ (10^−4^, 10^−2^)]. We do not model any pleiotropic effects. We select *IV_CIG_* and learn the causal graph skeleton and PAG at a threshold *α* = 10^−3^. We select *IV_G_* in a GWAS with a significance threshold of 10^−8^. We used the p-value thresholds in the range [10^−8^, 10^0^] to select significantly causal links from the MR tests. Lines show bootstrapped means, error bars indicate the bootstrapped 95% confidence interval of the mean (1000 samples), calculated across 20 replicates. Grey filled circles indicate the mean at a p-value threshold of 0.05. The dotted line indicates *TDR* = 0.95. A detailed description of all calculations and MR methods used is given in the Methods section.

**Fig. S4:**
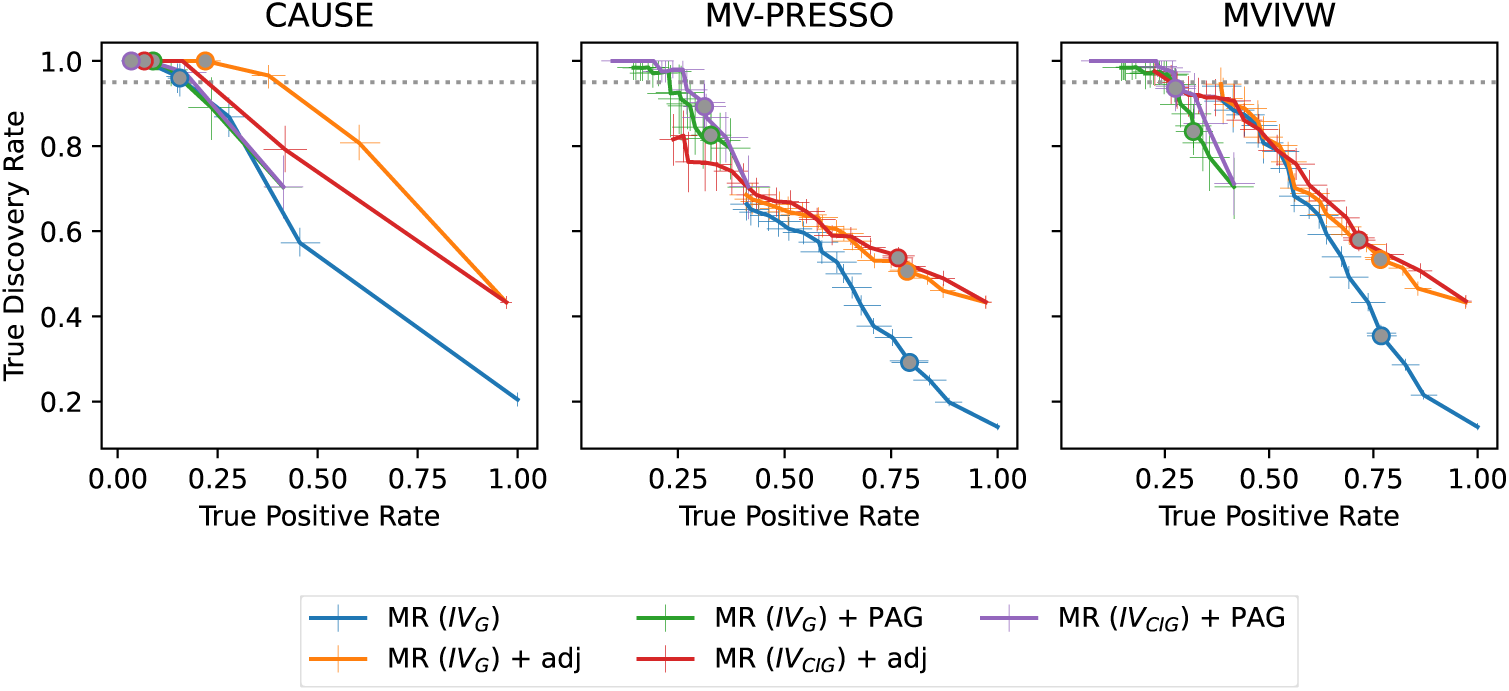
Causal inference performance of CI-GWAS with commonly used MR methods in simulation with strong LD and ten strongly correlated traits, using relaxed significance thresholds. Here we show the true discovery rate and true positive rate of the trait-trait causal link detection for a number of commonly used MR methods when used with GWAS selected instrumental variables (*IV_G_*), CI-GWAS selected instrumental variables (*IV_CIG_*), and with and without using the CI-GWAS skeleton (adj) or PAG as an additional filter for causal relations. In the simulations for this figure, we drew marker-trait effects from *U* [(−10^−2^, −10^−4^) ∪ (10^−4^, 10^−2^)], and a 0.2 chance for each marker to have a pleiotropic effect on an additional trait (*γ* = 0.2). We select *IV_CIG_* and learn the causal graph skeleton and PAG at a threshold *α* = 10^−2^. We select *IV_G_* in a GWAS with a significance threshold of 10^−4^. We used the p-value thresholds in the range [10^−8^, 10^0^] to select significantly causal links from the MR tests. Lines show bootstrapped means, error bars indicate the bootstrapped 95% confidence interval of the mean (1000 samples), calculated across 20 replicates. Grey filled circles indicate the mean at a p-value threshold of 0.05. The dotted line indicates *TDR* = 0.95. A detailed description of all calculations and MR methods used is given in the Methods section.

**Fig. S5:**
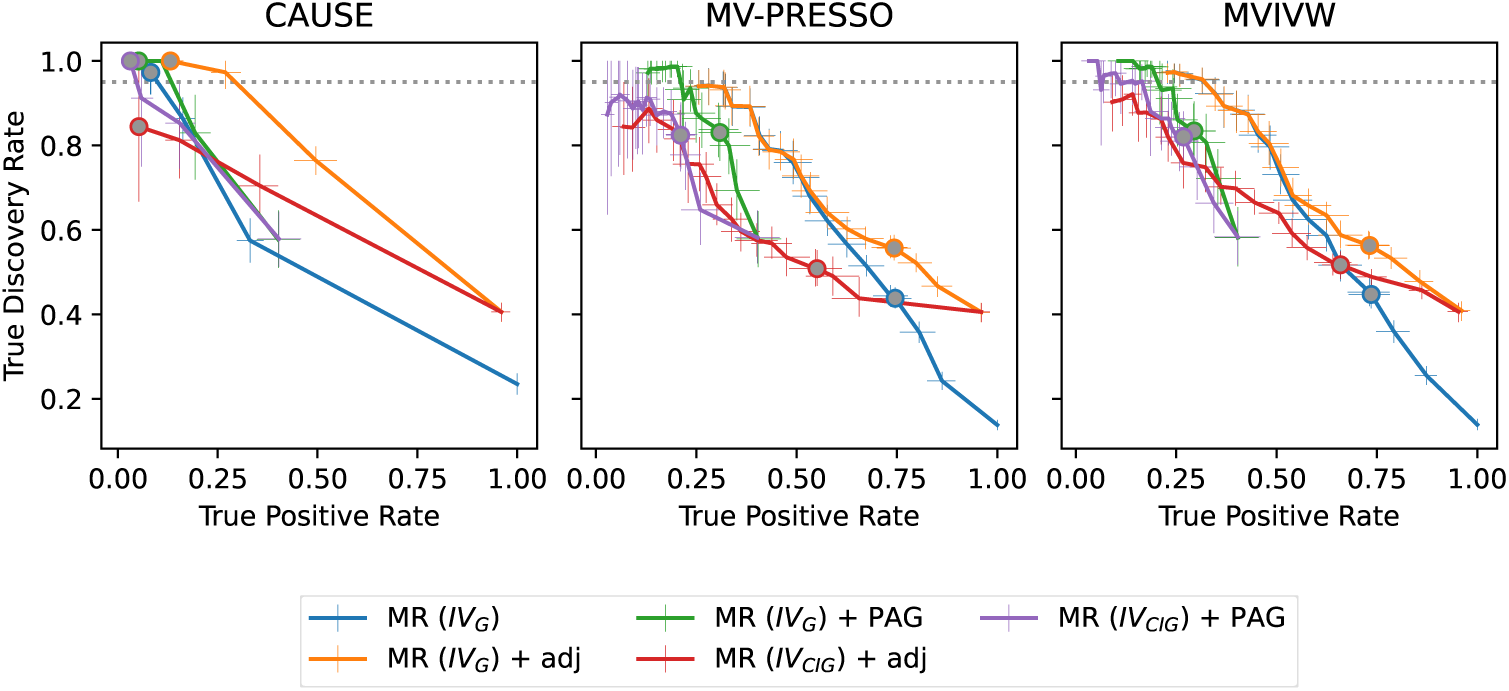
Causal inference performance of CI-GWAS with commonly used MR methods in simulation with strong LD and ten strongly correlated traits, using relaxed significance thresholds and without pleiotropy. Here we show the true discovery rate and true positive rate of the trait-trait causal link detection for a number of commonly used MR methods when used with GWAS selected instrumental variables (*IV_G_*), CI-GWAS selected instrumental variables (*IV_CIG_*), and with and without using the CI-GWAS skeleton (adj) or PAG as an additional filter for causal relations. In the simulations for this figure, we drew marker-trait effects from *U* [(−10^−2^, −10^−4^) ∪ (10^−4^, 10^−2^)]. We do not model any pleiotropic effects. We select *IV_CIG_* and learn the causal graph skeleton and PAG at a threshold *α* = 10^−2^. We select *IV_G_* in a GWAS with a significance threshold of 10^−4^. We used the p-value thresholds in the range [10^−8^, 10^0^] to select significantly causal links from the MR tests. Lines show bootstrapped means, error bars indicate the bootstrapped 95% confidence interval of the mean (1000 samples), calculated across 20 replicates. Grey filled circles indicate the mean at a p-value threshold of 0.05. The dotted line indicates *TDR* = 0.95. A detailed description of all calculations and MR methods used is given in the Methods section.

**Fig. S6:**
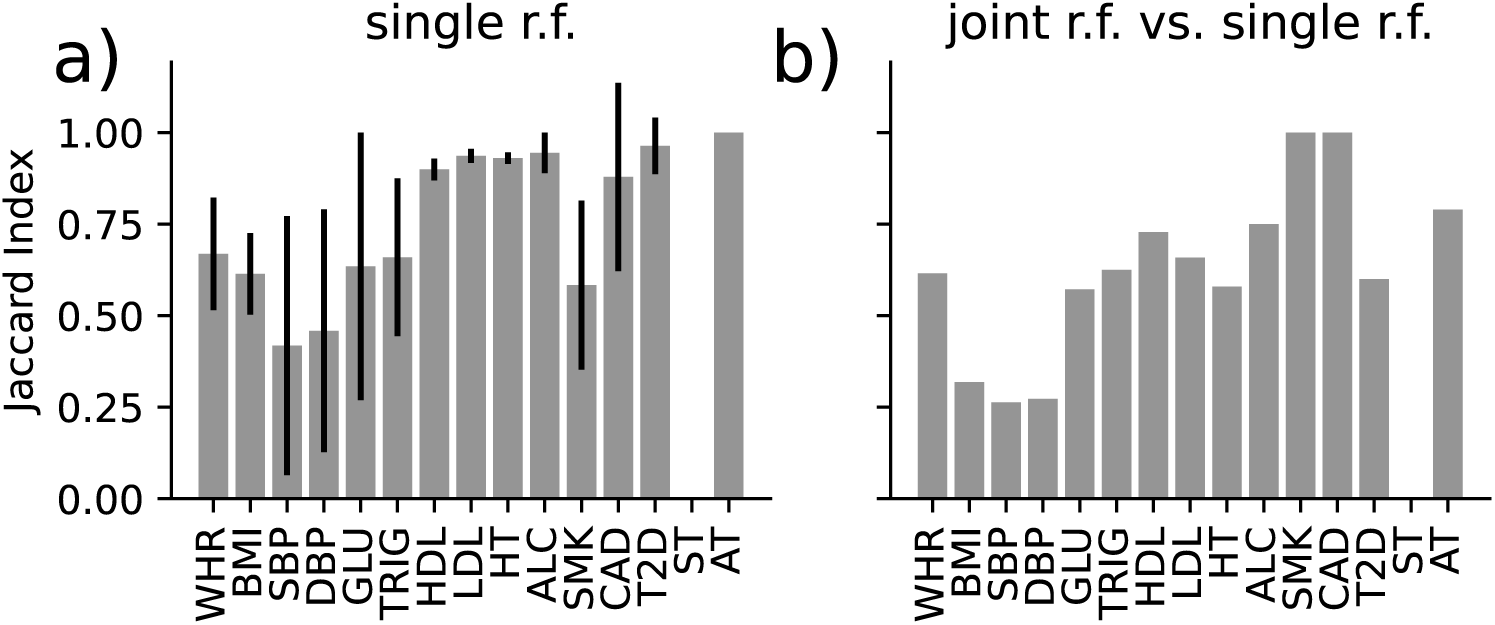
Similarities of SNP sets identified in causal inference in the UK Biobank with CI-GWAS. We analysed 36 risk factor (r.f.) - outcome relationships in the UK Biobank. We ran either pairs of a single risk factor and outcome (single r.f.), or all risk factors with a single outcome jointly (joint r.f.). (a): Jaccard Indices across all SNP sets identified for each trait in the single r.f. analyses. Errorbars show the 95% confidence interval. (b): Jaccard index of the intersection of the SNP sets identified the joint r.f. analyses and the intersection of the SNP sets identified in the single r.f. analyses, per trait.

**Fig. S7:**
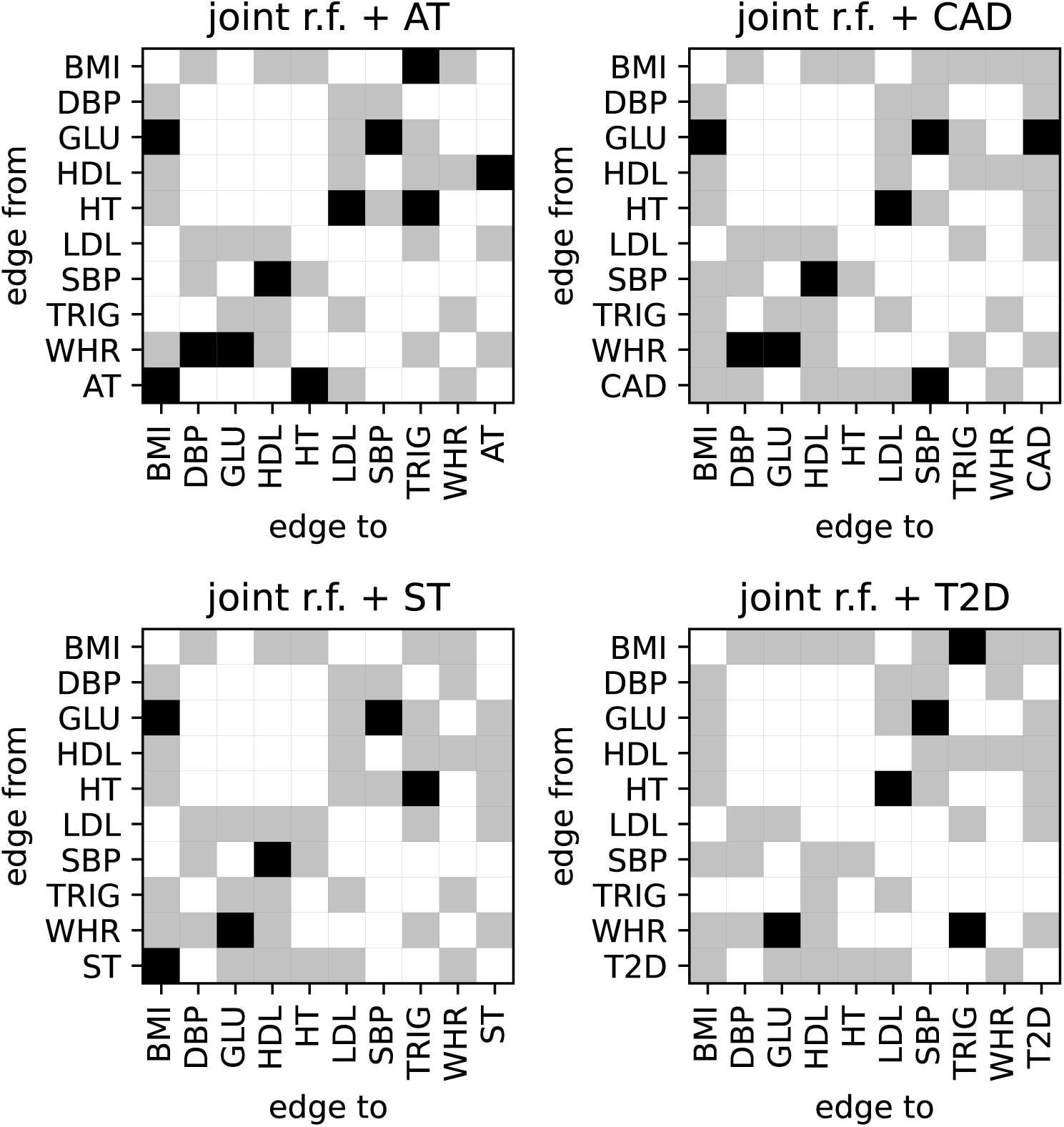
Partial ancestral graphs inferred in the UK Biobank with CI-GWAS. We analysed 36 risk factor (r.f.) - outcome relationships in the UK Biobank. We ran all risk factors with a single outcome jointly (joint r.f.). Black squares indicate unambigously directed edges. Gray squares indicate bidirected edges that indicate the presence of a latent confounder or a cyclical relationship that violates the inference assumptions. White squares indicate the non-adjacency.

**Fig. S8:**
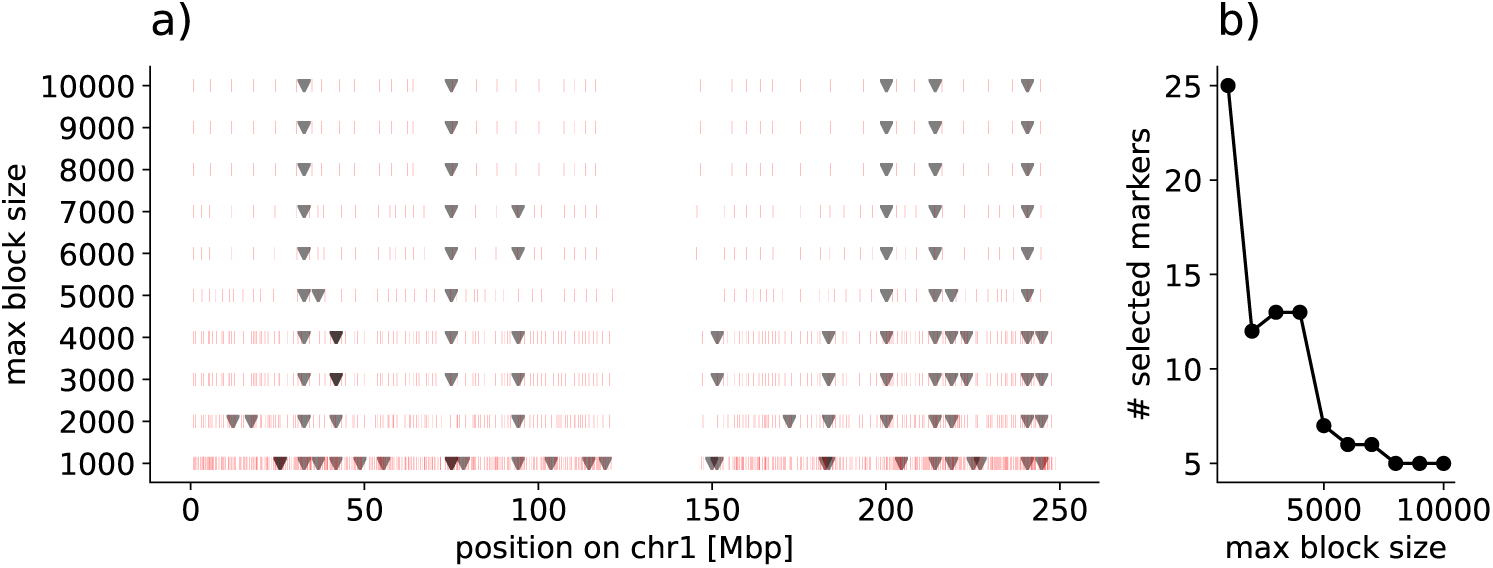
An example of the effect of maximum marker block size on the cuda-skeleton results. The results of the cuda-skeleton are shown across various sets of block definitions generated with model parameter max block size *s_max_* ranging from 1000 to 10000 markers. (a) gives the maximum block size used plotted against the positions of markers on chromosome 1. The position of markers selected as having a significant partial correlation with a trait is given by a triangle, and the marker block edges are given by red lines. (b) shows the total number of markers selected on chromosome 1, given the maximum block size used. These results were obtained from the cuda-skeleton with *α* = 10^−4^, using genomic data of chromosome 1 in the UK Biobank data and 17 traits (see Methods). Changing the marker block size either joins or separates markers that are correlated (those in linkage disequilibrium, LD). If two markers in the same block are in high LD, splitting them into two blocks can increase the partial correlation between markers within those blocks and traits, resulting in both markers being selected. Joining two neighbouring blocks, where markers previously separated are in LD, reduces marker associations. With increasing block size, fewer, more consistent markers are selected and thus we used the largest block size feasible for our GPU architecture.

---

**Table S1:**
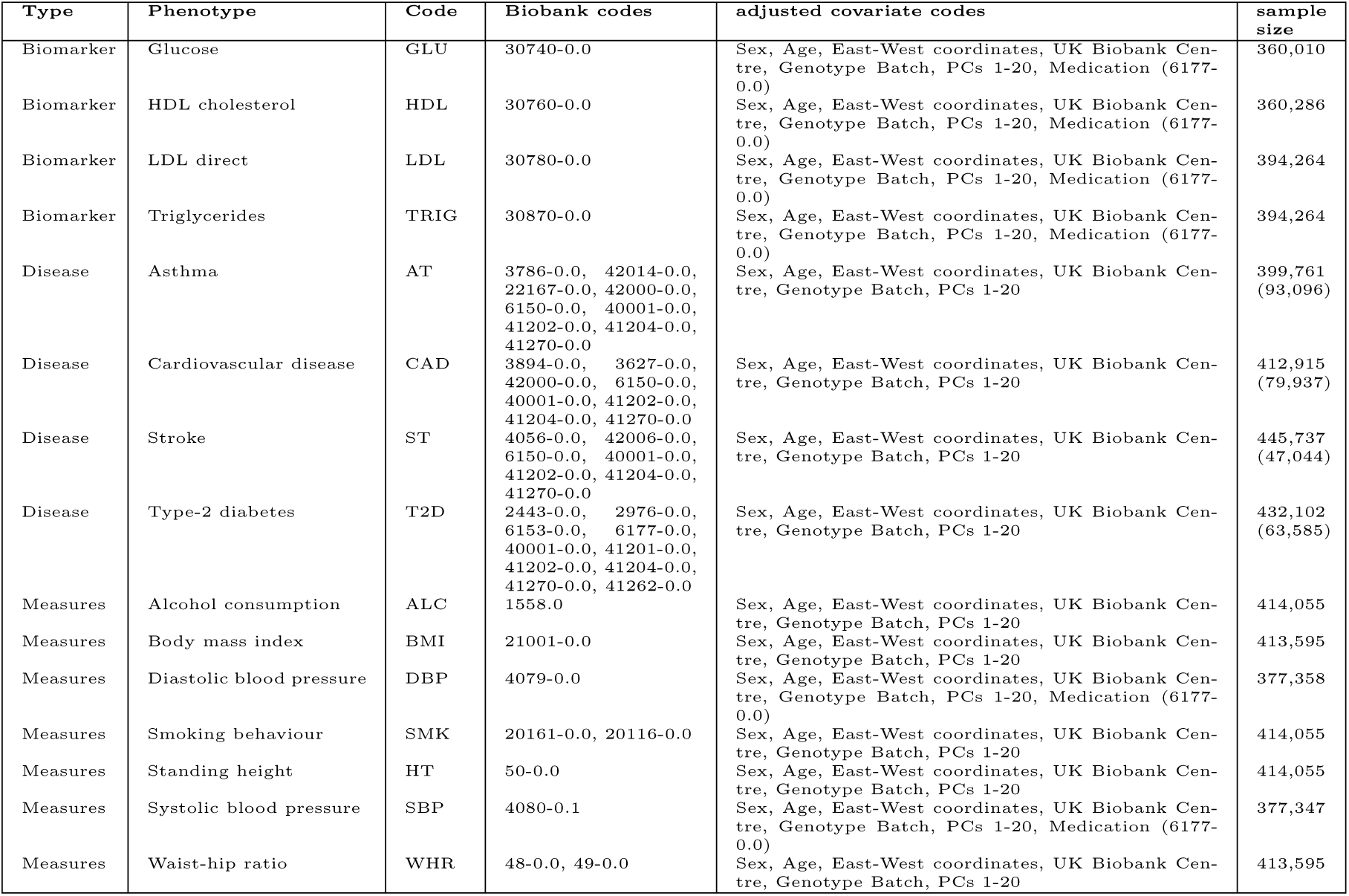
UK Biobank phenotypes used within the study. Columns show the type of trait, the name, the trait code used for the figures, the UK Biobank codes used to construct the trait values, the covariates adjusted for within the analysis, and the sample size (case numbers given in brackets).

